# Deciphering the association of spatial landscapes of DCIS with recurrence as invasive breast cancer

**DOI:** 10.64898/2026.09.25.754326

**Authors:** Nicola Graham, Arran K Turnbull, Carlos Martínez-Pérez, J Michael Dixon, Florent Petitprez, Takanori Kitamura

## Abstract

Stromal cell composition has been proposed as a potential biomarker in ductal carcinoma *in situ* (DCIS) for assessing the risk of subsequent recurrence or progression to invasive breast cancer (IBC). However, the characteristics of the tumor microenvironment within DCIS associated with these outcomes remain poorly understood. Here, we perform spatial transcriptomic analyses of 49 primary DCIS specimens from patients with no recurrence, DCIS recurrence, or recurrence as IBC. Cell mapping reveals spatially organized stromal cell gradients within DCIS, with enrichment of TREM2^+^ macrophages and myofibroblastic cancer-associated fibroblasts (myCAFs) in tumor-proximal niches. Notably, the abundance of TREM2^+^ macrophages near tumor nests is associated with subsequent recurrence as IBC. Moreover, ligand-receptor pair analysis identifies GRN–SORT1, SPP1–CD44, and LGALS9– CD44 interactions as spatially plausible signaling interactions between TREM2⁺ macrophages and epithelial cells. Finally, these findings are validated at the protein level by multiplex immunostaining of serial sections. This study characterizes the spatial landscape of the DCIS tumor microenvironment associated with subsequent recurrence, providing a basis for developing biomarkers of disease progression and identifying potential therapeutic targets.

## Introduction

Ductal carcinoma *in situ* (DCIS), the earliest stage of breast cancer, accounts for up to 25% of all breast cancer diagnoses, with over 60,000 new cases in the USA and more than 7,000 in the UK diagnosed annually ^[1]^. DCIS is a non-invasive breast neoplasm that originates from the mammary ductal epithelium and remains confined within the ducts. However, some DCIS lesions can progress to invasive breast cancer (IBC), which may subsequently develop into life-threatening metastatic disease ^[2,3]^. Consequently, conventional treatment of DCIS includes mastectomy or breast-conserving surgery, with radiotherapy and adjuvant endocrine therapy recommended in selected cases. However, in cohorts of patients with untreated or unresected DCIS, approximately 67-88% of patients remained free from invasive disease during follow-up ^[4–6]^, suggesting that a substantial proportion of DCIS may not progress to IBC. This raises concerns regarding potential overtreatment, particularly among patients with low-risk disease, as conventional treatment can substantially affect quality of life ^[1]^. Conversely, despite treatment, a proportion of patients with DCIS subsequently develop recurrent disease, which may present as recurrent DCIS or as IBC ^[7]^. Thus, the clinical challenge lies in avoiding unnecessary treatment of indolent DCIS while ensuring that patients at increased risk of recurrence or progression to IBC receive appropriate treatment. Nevertheless, it remains challenging to accurately distinguish indolent DCIS from lesions with a high risk of recurrence or progression to IBC. Therefore, there is an unmet clinical need for biomarkers that can identify, at the time of initial diagnosis, DCIS patients at increased risk of subsequent recurrence or progression to IBC, thereby enabling more accurate risk stratification and personalized treatment.

To identify biomarkers predictive of DCIS progression, previous studies have investigated molecular differences in tumor epithelial cells between the normal breast, DCIS, and IBC tissues. Although the transcriptomic features of epithelial cells in DCIS differ substantially from those in normal breast tissue, gene expression profiles of epithelial cells in DCIS and IBC have been reported to be relatively comparable ^[3]^. This suggests that differences in tumor epithelial cell-intrinsic transcriptional programs alone may not fully explain the transition from DCIS to IBC. Consequently, recent studies have increasingly focused on alterations in the surrounding tumor microenvironment (TME) that may contribute to DCIS progression and/or recurrence. Several immunohistochemical studies have demonstrated increased numbers of immune cell populations, including regulatory T (Treg) cells and macrophages, as well as increased expression of immune checkpoints such as PD-L1, in IBC compared with DCIS ^[8–12]^. These findings suggest that the establishment of an immunosuppressive TME may contribute to the progression of DCIS to IBC. More recently, single-cell and bulk transcriptomic studies have further characterized the cellular and molecular composition of the DCIS microenvironment ^[13,14]^. However, the spatial organization of these cellular populations within the DCIS microenvironment and their association with subsequent disease recurrence or progression to IBC remain poorly understood, despite findings from advanced breast cancer that spatially distinct TME niches are associated with clinical outcomes ^[15]^.

To address these knowledge gaps, we perform spatial transcriptomic profiling using the CosMx platform with a 6K-gene panel in a cohort of 49 primary DCIS samples (21 cases without recurrence, 14 with subsequent DCIS recurrence, and 14 with IBC recurrence). Additionally, we perform single-nuclei RNA sequencing (snRNA-seq) on an independent cohort of DCIS and IBC (4 low-grade DCIS, 4 high-grade DCIS and 4 IBC) and integrate the results with the spatial data set. This approach enables the delineation of a high-resolution cell atlas and more comprehensive gene expression profiling of individual cells, revealing the following observations; (1) gradient-like spatial distributions of macrophage and cancer-associated fibroblast (CAF) subpopulations within the DCIS TME; (2) an association between TREM2^+^ macrophage abundance within the tumor-proximal niche, myofibroblastic CAF (myCAF) localization, and subsequent IBC recurrence; (3) spatially plausible ligand–receptor interactions between TREM2^+^ macrophage populations and epithelial cells in DCIS; and (4) an association between TREM2^+^ macrophage abundance within spatially distinct tumor-proximal niches and recurrence status that was recapitulated by immunostaining. Together, these findings identify spatially distinct TME niches in DCIS and provide potential biomarkers and therapeutic targets for risk stratification and prevention of invasive recurrence.

## Results

### Transcriptionally distinct cell types in DCIS revealed by single-nuclei RNA-sequencing

To characterize the cellular composition of DCIS, we performed snRNA-seq on 8 DCIS samples, including 4 low/intermediate-grade and 4 high-grade cases, alongside 4 IBC samples (**Fig. 1a**, **Supplementary Table 1**). Following data processing, quality filtering, and curation, a total of 179,195 cells were retained for downstream analysis. Based on the expression of key marker genes, each cell was assigned to one of the main lineages; epithelial cell, myeloid cell, lymphocyte, fibroblast and pericyte, endothelial cell, or adipocyte (**Fig. 1b**, **Supplementary Fig. 1a**). Within each lineage, cells were re-clustered and assigned to cell subset identities.

In the epithelial compartment, cells were segregated to two subsets based on their ploidy estimated by the Copy number Karyotyping of Aneuploid Tumors (CopyKAT) approach ^[16]^. Epithelial cells classified as diploid were annotated as normal epithelial (N-Epi) cells, while those classified as aneuploid were annotated as tumor epithelial (T-Epi) cells. Consistent with a previous report demonstrating the presence of copy number abnormalities in DCIS tissues ^[17]^, DCIS cases in our dataset contained both N-Epi and T-Epi cells, with N-Epi cells predominating in DCIS and T-Epi cells enriched in IBC (**Fig. 1c**).

Based on the expression of lineage marker genes, myeloid cells were subdivided into four macrophage populations (TREM2^+^, FOLR2^+^, SPP1^+^, and IL4I1^+^ macrophage), classical monocytes, three dendritic cell (DC) populations (conventional DC 1; cDC1, conventional DC 2; cDC2, and CCR7^+^ DC), and mast cells (**Fig. 1d**,**e**). Lymphocytes were subdivided into three T cell populations (CD8^+^ T cells, CD4^+^ T cells, and Treg cells), NK cells, two B cell subsets (naïve and memory B), and plasma cells (**Fig. 1d,f**). The fibroblast-pericyte lineage was split into three CAF populations (myCAF, inflammatory CAF [iCAF], and CD34^+^ CAF), myoepithelial cells, and pericytes (**Fig. 1d,g**). Endothelial cells were divided into two subpopulations: blood vascular endothelial cells and lymphatic endothelial cells (**Fig. 1d**, **Supplementary Fig. 1b**).

While all 26 cell populations were identified in DCIS, not all cell clusters were equally represented across samples of different grades. Notably, several clusters were enriched in either low- or high-grade DCIS samples, whereas others were predominantly found in IBC samples (**Supplementary Fig. 1c,d**). These findings suggest that DCIS may exhibit distinct tumor microenvironment profiles compared with IBC and therefore requires specific analysis.

**Fig. 1.**
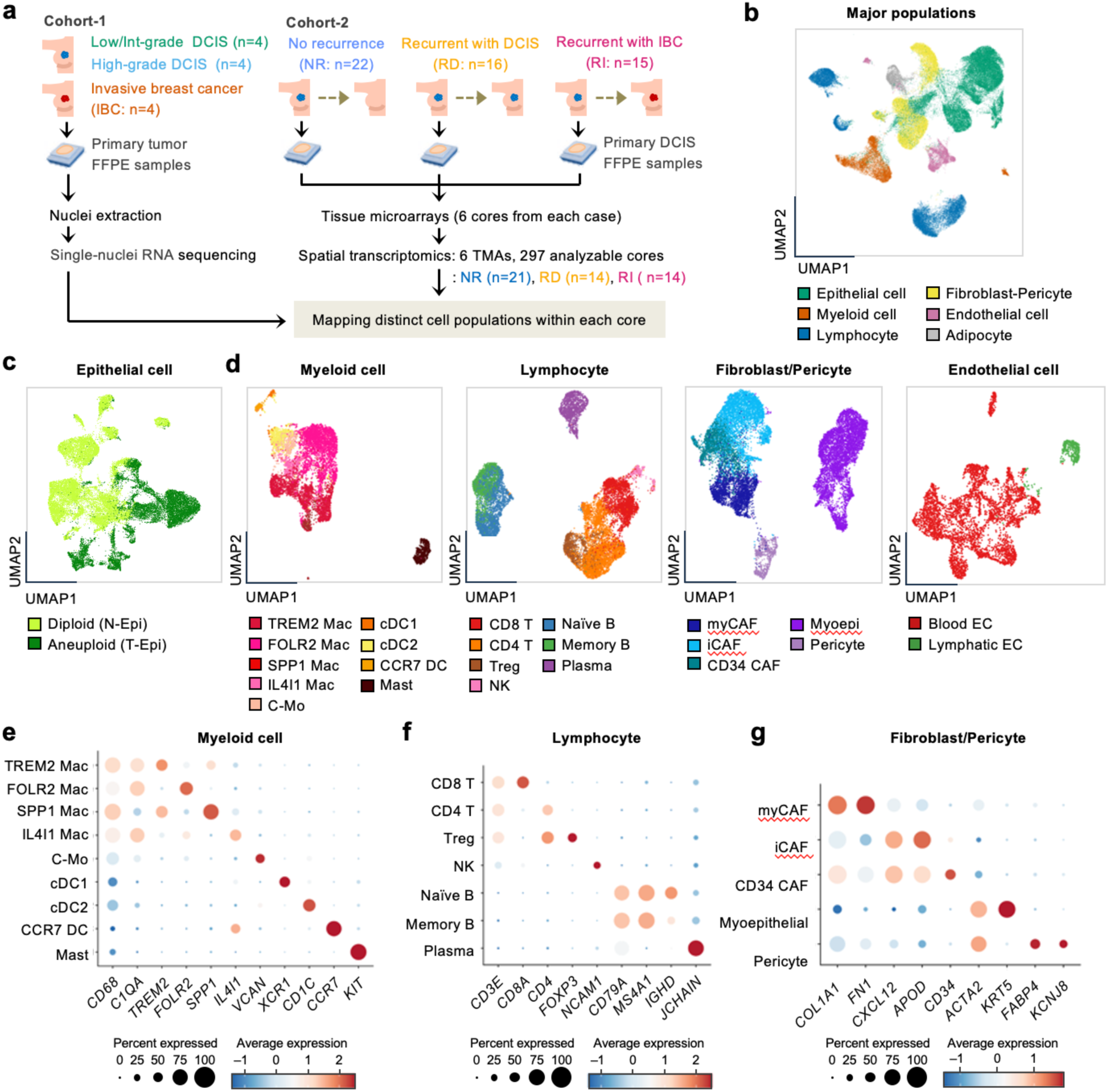
Identification of cell types within the DCIS tissues. **a.** Study overview. FFPE tissues of 4 low/intermediate-grade DCIS, 4 high-grade DCIS, and 4 IBC cases were analyzed by snRNA-seq. A DCIS-derived reference matrix was used for cell type annotation in spatial transcriptomics. Six TMAs comprising primary DCIS tissues from an independent cohort were analyzed by CosMx SMI (6,000-plex panel), followed by cell segmentation and cell type annotation. Cases were classified as recurrence with IBC (RI), recurrence with DCIS but not IBC (RD), or no recurrence (NR). After quality control, 297 cores from 49 cases were retained. **b.** UMAP plot of snRNA-seq data showing major cell lineages. **c.** UMAP plots showing two epithelial cell subsets defined based on their ploidy. **d.** UMAP plots showing subpopulations of myeloid cells, lymphocytes, fibroblasts/pericytes, and endothelial cells identified by lineage-specific re-clustering. **e-g.** The expression of lineage marker genes in myeloid (e), lymphocyte (f), and fibroblast/pericyte (g) subpopulations. Circle size and color indicate the percentage of cells expressing each gene and its average expression level, respectively. FFPE, formalin-fixed paraffin-embedded; DCIS, ductal carcinoma in situ; IBC, invasive breast cancer; snRNA-seq, single-nuclei RNA sequencing; TMA, tissue microarray; Mac, macrophage; C-Mo, classical monocyte; DC, dendritic cell; Treg, regulatory T cell; NK, natural killer cell; CAF, cancer-associated fibroblast.

### Spatial distribution of cell types in DCIS lesions revealed by spatial transcriptomics

To visualize the spatial distribution of cell types in the microenvironment of DCIS, we performed spatial transcriptomics on a retrospective case-control cohort of DCIS samples (**Fig. 1a**). The cohort included the primary DCIS tissues from patients who recurred with IBC (recurrent IBC (RI), n = 15) or DCIS but not IBC (recurrent DCIS (RD), n = 16) following surgery. Additionally, we selected the tissues from patients who remained recurrence-free until the end of follow-up up to 11 years (non-recurrence (NR), n = 22). From each case, 6 cores containing tumor ducts and surrounding stroma were selected and embedded across 6 separate tissue microarray (TMA) blocks. Spatial transcriptomics was then performed using Nanostring CosMx 6K panel. For quality control, we retained only regions of interest (ROIs) with more than 200 transcripts per cell. Immunostaining images were also evaluated, and cores or ROIs lacking pan-cytokeratin-positive epithelial cells or showing tissue detachment were excluded. Following quality control and filtering, a total of 175 cores and 689 ROIs from 21 NR, 14 RD, and 14 RI cases were retained for downstream analysis (**Fig. 1a**, **Supplementary Table 3**). All groups included a comparable proportion of high-grade (Grade 3) and low/intermediate-grade (Grade 1 and 2) cases (**Supplementary Table 2**). Following data normalization, cell types were annotated at the single-cell level using supervised classification based on a reference matrix comprising cell type-specific gene expression profiles derived from the DCIS cases in the snRNA-seq dataset in Figure 1. All cells were assigned to one of the 16 major cell types (**Fig. 2a**). Consistent with the snRNA-seq analysis, epithelial cells, macrophages, dendritic cells, B cells, fibroblasts, and endothelial cells were further classified into their respective subpopulations (**Fig. 2b**). No substantial differences in data quality were observed across TMAs or cell types (**Supplementary Fig. 2**).

In our dataset, most epithelial cells were annotated as normal epithelium, with a smaller proportion classified as tumor epithelium (**Fig. 2c**). Within the stromal compartment, CAFs and myoepithelial cells were the most abundant non-immune cell types. The immune cell compartment was dominated with CD4^+^ T cells, dendritic cells, monocytes/macrophages, and mast cells, whereas CD8^+^ T cells, regulatory T cells, B cells, and NK cells were comparatively rare (**Fig. 2c**). Among the monocyte/macrophage population, TREM2^+^, FOLR2^+^, and IL4I1^+^ macrophages represented the predominant subpopulations, while cDC2 was the predominant DC subset. Within the CAF compartment, iCAFs and myCAFs represented the predominant subpopulations, whereas CD34+ CAFs were rarely observed Within the endothelial compartment, blood endothelial cells were predominant, whilst lymphatic cells were relatively rare (**Fig. 2d**).

### Epithelial cells composing the tumor nest in DCIS

Across patient groups, including non-recurrence (NR), recurrent DCIS (RD), and recurrent IBC (RI), tumor nests comprised both N-Epi and T-Epi cells (**Supplementary Fig. 3a**). As recurrent DCIS cases have been reported to exhibit a higher burden of copy number aberrations compared with non-recurrent cases ^[18]^, we investigated whether the proportion of T-Epi cells within tumor nests differed among patient groups. Tumor nests were defined as spatially contiguous epithelial clusters containing at least six contacting epithelial cells (**Fig. 2e**). Enclosed non-cellular spaces within these defined tumor nests were assigned as ductal (luminal) cavities. Smaller epithelial fragments and isolated epithelial cells were excluded from analysis, as they lacked sufficient spatial organization to represent coherent tumor structures. Comparison of the T-Epi fraction within the tumor nests revealed no significant differences between recurrence and non-recurrence groups (**Fig. 2f**). We further examined transcriptional differences among epithelial cells comprising tumor nests across patient groups. However, no significant differences were identified in either N-Epi or T-Epi cells in any comparisons (adjusted *P* > 0.6; data not shown). Together, these findings indicate that DCIS recurrence status is not associated with measurable differences in the cellular composition or transcriptional state of epithelial cells, prompting further characterization of the stromal compartment.

**Fig. 2.**
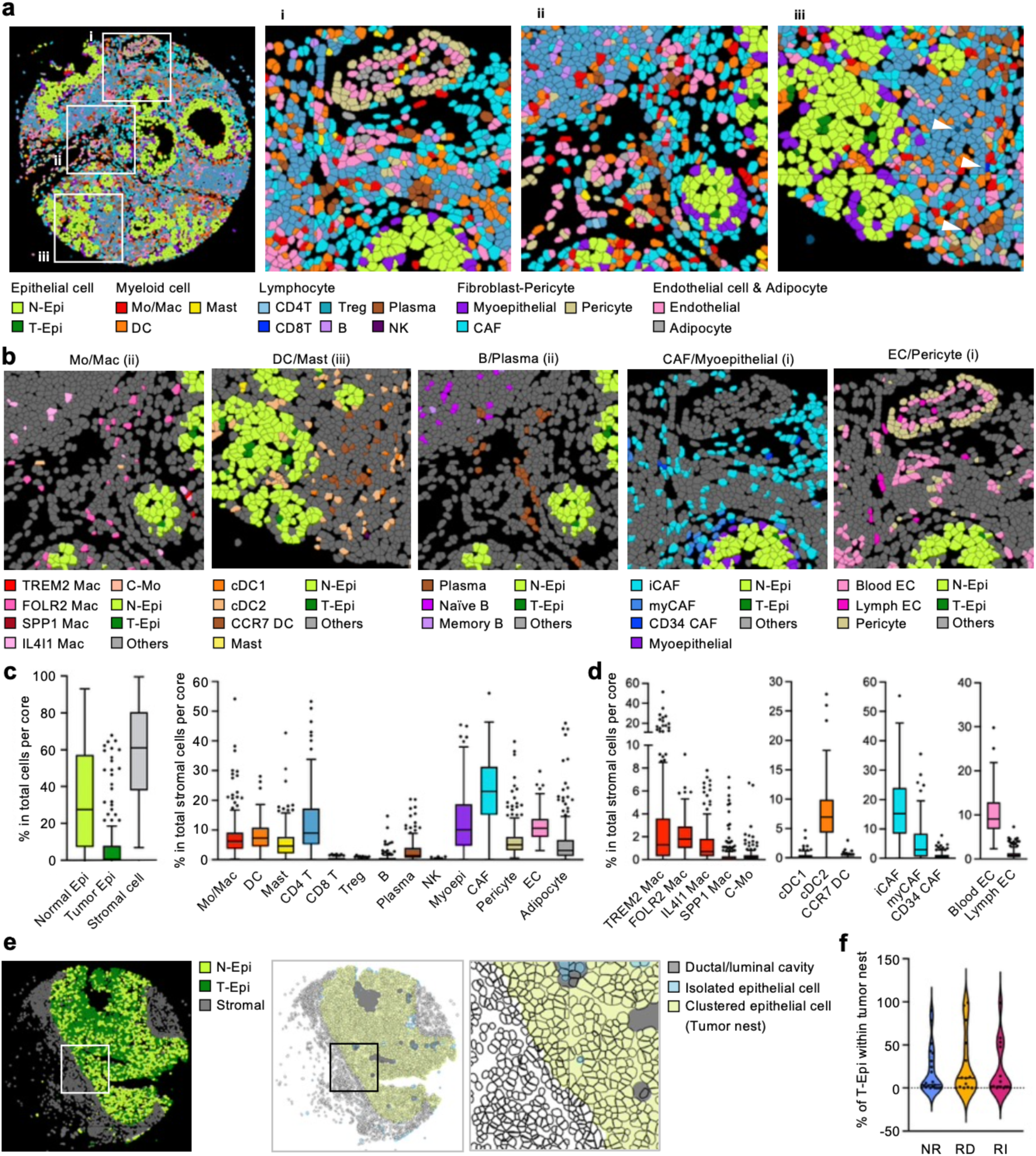
Spatial mapping of cell populations within the DCIS tumor microenvironment. **a.** Representative spatial transcriptomic image of a core after annotation of major cell lineages. High-magnification images of selected areas (i–iii) are also shown. **b.** High-magnification images of selected areas (i–iii) showing subpopulations defined in Fig. 1. **c.** Tukey box plots showing the proportions of epithelial and stromal cells among all cells (left) and major lineages among stromal cells (right) per core. Boxes represent the 25th–75th percentiles, with the median shown as a horizontal line. Whiskers extend to the most extreme data points within 1.5 × the interquartile range of the respective quartile. Values beyond the whiskers are plotted individually as outliers. **d.** Tukey box plots showing the proportions of Mac, DC, CAF, and EC subpopulations among total stromal cells per core. **e.** Representative images of a core showing the distribution of epithelial and stromal cells (left) and defined tumor nests, isolated epithelial cells, and luminal cavities (middle). High-magnification images of the selected area are shown on the right. **f.** Violin plot showing the proportion of T-Epi cells among epithelial cells within tumor nests across patient groups (NR, RD, and RI). Each dot represents one case. No statistical significances were detected by Kruskal–Wallis tests followed by Dunn’s multiple-comparison test. N-Epi, normal epithelial cell; T-Epi, tumor epithelial cell; Mac, macrophage; C-Mo, classical monocyte; DC, dendritic cell; Treg, regulatory T cell; NK, natural killer cell; CAF, cancer-associated fibroblast; EC, endothelial cell; NR, no recurrence; RD, recurrent DCIS; RI, recurrent IBC.

### Stromal cells organize on a gradient around tumor cores

To investigate stromal features associated with recurrence, we categorized the TME into five spatial niches according to their relationship to tumor nests (**Fig. 3a**). The inside-nest/contact with epithelium (IC) niche comprised stromal cells located within tumor nests and in direct contact with epithelial cells, whereas the inside-nest/duct (ID) niche comprised stromal cells located within ductal (luminal) cavities inside tumor nests. The outside-nest/contact with epithelium (OC) niche comprised stromal cells immediately adjacent to tumor nests, the outside-nest/near epithelium (ON) niche comprised stromal cells within the four-cell-distance peri-tumoral region, and the outside-nest/far from epithelium (OF) niche comprised stromal cells located distal to tumor nests. Although tumor nests comprised both N-Epi and T-Epi cells, the proportion of T-Epi cells within tumor nests was not strongly correlated with the abundance of any stromal cell subset (**Supplementary Fig. 3b**). The boundary of the ON niche was defined based on a previous study showing that cytokines released by immune cells can diffuse over distances of approximately 30–40 µm ^[19]^, suggesting that cells within four-cell-diameters may be exposed to immune cell-derived signals. This spatial framework enabled quantitative comparison of stromal cell composition across distinct tumor-associated microenvironments.

We then determined the distribution of non-immune stromal cells across spatial niches. As expected, myoepithelial cells, which typically form a thin layer beneath the mammary glandular epithelium, were predominantly localized near epithelial structures, particularly tumor-proximal IC and OC niches. In contrast, iCAFs and CD34^+^ CAFs were enriched in the tumor-distal OF niche. Although myCAFs were also present in the OF niche, they showed greater enrichment in the tumor-proximal ON niche (**Fig. 3b**). These findings demonstrate a spatial gradient of epithelial/fibroblast-associated populations, transitioning from myoepithelial cells near tumor nests to myCAFs and subsequently to iCAFs/CD34+ CAFs at increasing distances from the tumor epithelium. Lymphatic endothelial cells and adipocytes were predominantly localized to the tumor-distal OF niche, whereas pericytes and blood endothelial cells were distributed across all spatial niches (**Supplementary Fig. 4a**). We performed similar analysis on the immune cell compartment and found that macrophage subpopulations also exhibited distinct spatial gradients. Namely, TREM2^+^ macrophages were predominantly localized within the intra-ductal ID niche and tumor-proximal IC and OC niches. In contrast, FOLR2^+^ macrophages were rarely observed within tumor nests and progressively increased in abundance with increasing distance from tumor nests, reaching their highest representation in the tumor-distal OF niche (**Fig. 3c**). IL4I1^+^ macrophages were also predominantly localized outside tumor nests, with comparable distributions between the tumor-proximal OC/ON niches and the tumor-distal OF niche. SPP1^+^ macrophages and classical monocytes were exclusively detected in the OF niche, although their proportions were considerably lower than those of other macrophage subpopulations. The remaining immune cell populations were either predominantly localized to the tumor-distal OF niche (cDC1, CD8^+^ T, Treg, B, plasma, and NK cells) or distributed across multiple niches (cDC2, mast, and CD4^+^ T cells) (**Supplementary Fig. 4b**).

**Fig. 3.**
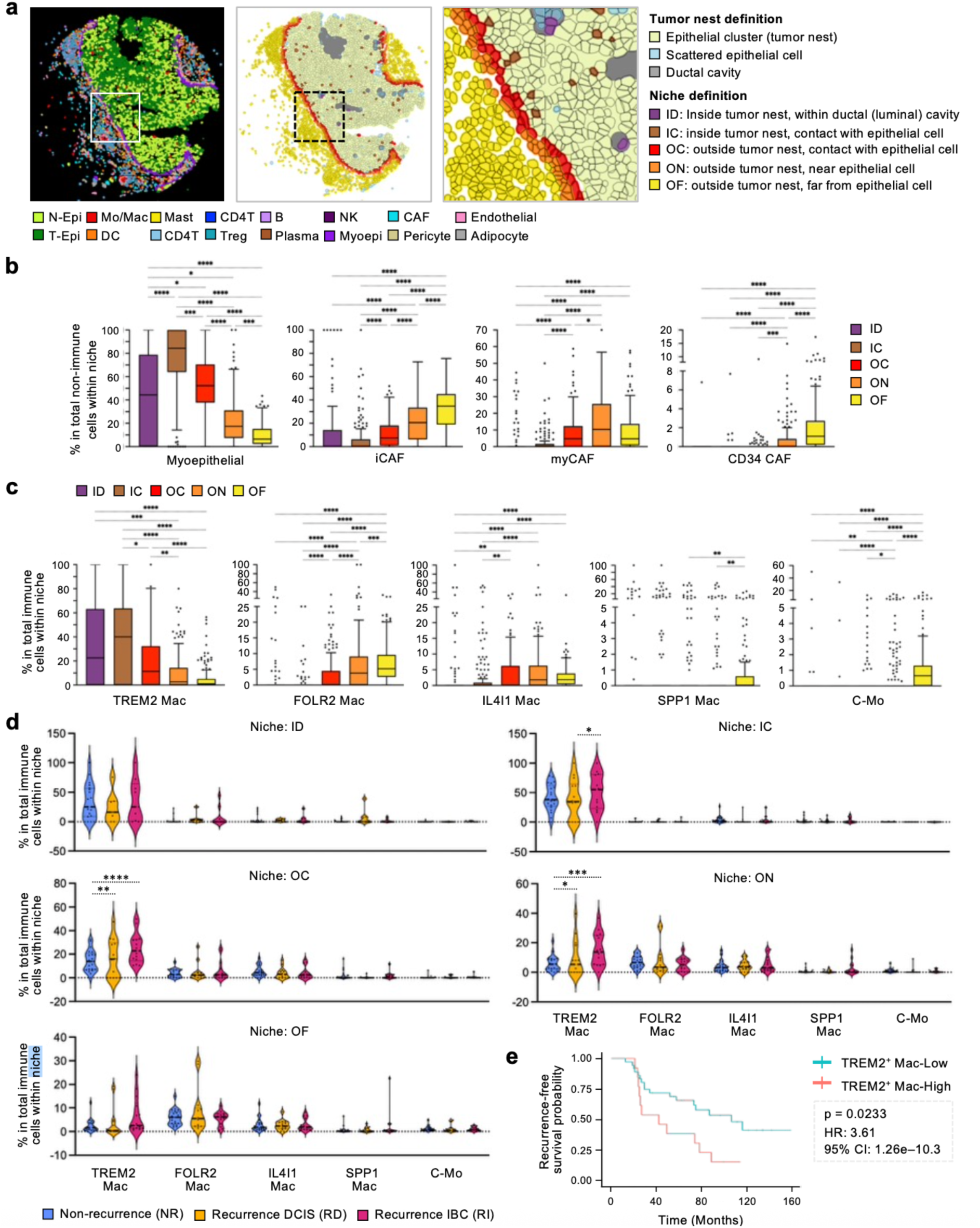
Spatially distinct localization of stromal cell subsets in tumor-associated niches. **a.** Representative images of a core shown in Fig. 2e displaying the distribution of major cell lineages (left) and spatially distinct niches defined according to their relationship to tumor nests (middle). High-magnification image of the selected area is shown on the right. **b.** Tukey box plots showing the proportions of myoepithelial cells and fibroblast subsets among non-immune stromal cells within each niche. **c.** Tukey box plots showing the proportions of macrophage subsets and monocytes among immune cells within each niche. Boxes represent the 25th–75th percentiles, with the median shown as a horizontal line. Whiskers extend to the most extreme data points within 1.5 × the interquartile range of the respective quartile. Values beyond the whiskers are plotted individually as outliers. Statistical differences across niches were assessed using Kruskal–Wallis tests followed by Dunn’s multiple-comparison test (\**p* < 0.05, \*\**p* < 0.01, \*\*\**p* < 0.001, and \*\*\*\**p* < 0.0001). **d.** Violin plots showing the proportions of macrophage subsets and monocytes among immune cells within each niche across patient groups (NR, RD, and RI). Each dot represents one case. Statistical differences among patient groups were assessed by two-way ANOVA followed by Tukey’s multiple-comparisons test (\**p* < 0.05, \*\**p* < 0.01, \*\*\**p* < 0.001, and \**p* < 0.0001). **e.** Kaplan-Meier curve showing recurrence-free survival. Groups were defined by the fraction of TREM2^+^ macrophages in the OC/ON niche: TREM2^+^ Mac-High (upper quartile, ≥25%) vs. TREM2^+^ Mac-Low (lower quartiles). p-value was assessed using a logrank test. Hazard ratio and 95% confidence interval were assessed using a Cox proportional hazard regression. N-Epi, normal epithelial cell; T-Epi, tumor epithelial cell; Mac, macrophage; C-Mo, classical monocyte; DC, dendritic cell; Treg, regulatory T cell; NK, natural killer cell; CAF, cancer-associated fibroblast; EC, endothelial cell; ID, inside-nest/duct; IC, inside-nest/contact with epithelium; OC, outside-nest/contact with epithelium; ON, outside-nest/near epithelium; OF, outside-nest/far from epithelium; NR, no recurrence; RD, recurrent DCIS; RI, recurrent IBC; HR, hazard ratio; CI, confidence interval.

### Tumor-proximal TREM2^+^ macrophages associate with DCIS recurrence as IBC

We next investigated whether the spatially distinct distributions of stromal cells were associated with DCIS recurrence status. For each niche, we quantified the fractions of immune and non-immune cell populations within their respective compartments and compared them across patient groups; NR, RD, and RI. Within the non-immune cell compartment, the abundance of cell populations, including myoepithelial cells and CAFs, were not changed among the groups (**Supplementary Fig. 5a**). Within the immune cell compartment, the proportion of TREM2^+^ macrophages did not differ significantly between the groups in the intra-cavity ID niche (**Fig. 3d**). In the intra-tumoral IC niche, TREM2^+^ macrophages were significantly more abundant in RI than in RD cases, whereas their abundance did not differ significantly between RI and NR cases. Moving towards the tumor-proximal OC and ON niches, TREM2^+^ macrophages were markedly more abundant in RI cases than in NR cases, with a similar but less pronounced increase in RD cases. In the tumor-distal OF niche, their abundance was comparable between the groups. These niche-dependent, recurrence-associated changes were not observed in other macrophage subpopulations. Although mast cell fractions in the ON and OF niches were lower in RD than in NR, and cDC2 and plasma cell fractions in the ON and/or OF niches were higher in RD than in RI, no significant differences were detected between RI and NR (**Supplementary Fig. 5b**). These findings identified tumor-proximal accumulation of TREM2^+^ macrophages as the most prominent spatial immune feature associated with DCIS recurrence, whereas other stromal cell populations showed limited or inconsistent recurrence-associated changes.

We further investigated whether the abundance of TREM2^+^ macrophages in the tumor-proximal OC/ON niches influenced the time to recurrence. The cohort was stratified into ‘high’ and ‘low’ groups based on the fraction of TREM2^+^ macrophages in the OC/ON niches, using the third quartile as the cutoff. Kaplan-Meier curves were then generated to analyze the time to recurrence for each group (**Fig. 3e**). A high fraction of TREM2^+^ macrophages was associated with a significantly shorter time to recurrence (hazard ratio 3.61, 95% confidence interval 1.26-10.3, logrank test p-value: 0.0233). Collectively, these results suggest that the localized enrichment of TREM2^+^ macrophages in tumor-proximal niches is a strong parameter for predicting the aggressiveness of recurrent disease.

### Colocalization of tumor-proximal TREM2^+^ macrophages and myCAFs influence DCIS recurrence

Next, we examined the stromal cell populations adjacent to TREM2^+^ macrophages within tumor-proximal niches and assessed whether their neighboring patterns differed across patient groups. To this end, we identified individual TREM2^+^ macrophages within OC/ON niches as reference cells and quantified stromal cells located within four-cell-diameters of each reference cell, defining a local environment (hereafter referred to as “milieu”) in which cells may receive signals from TREM2^+^ macrophages ^[19]^ (**Fig. 4a**). The most abundant cell types within this milieu were myoepithelial cells and TREM2^+^ macrophages, followed by blood ECs, cDC2, myCAFs, CD4^+^ T cells, and pericytes (**Fig. 4b**). Other stromal cells were rarely observed within this milieu.

**Fig. 4.**
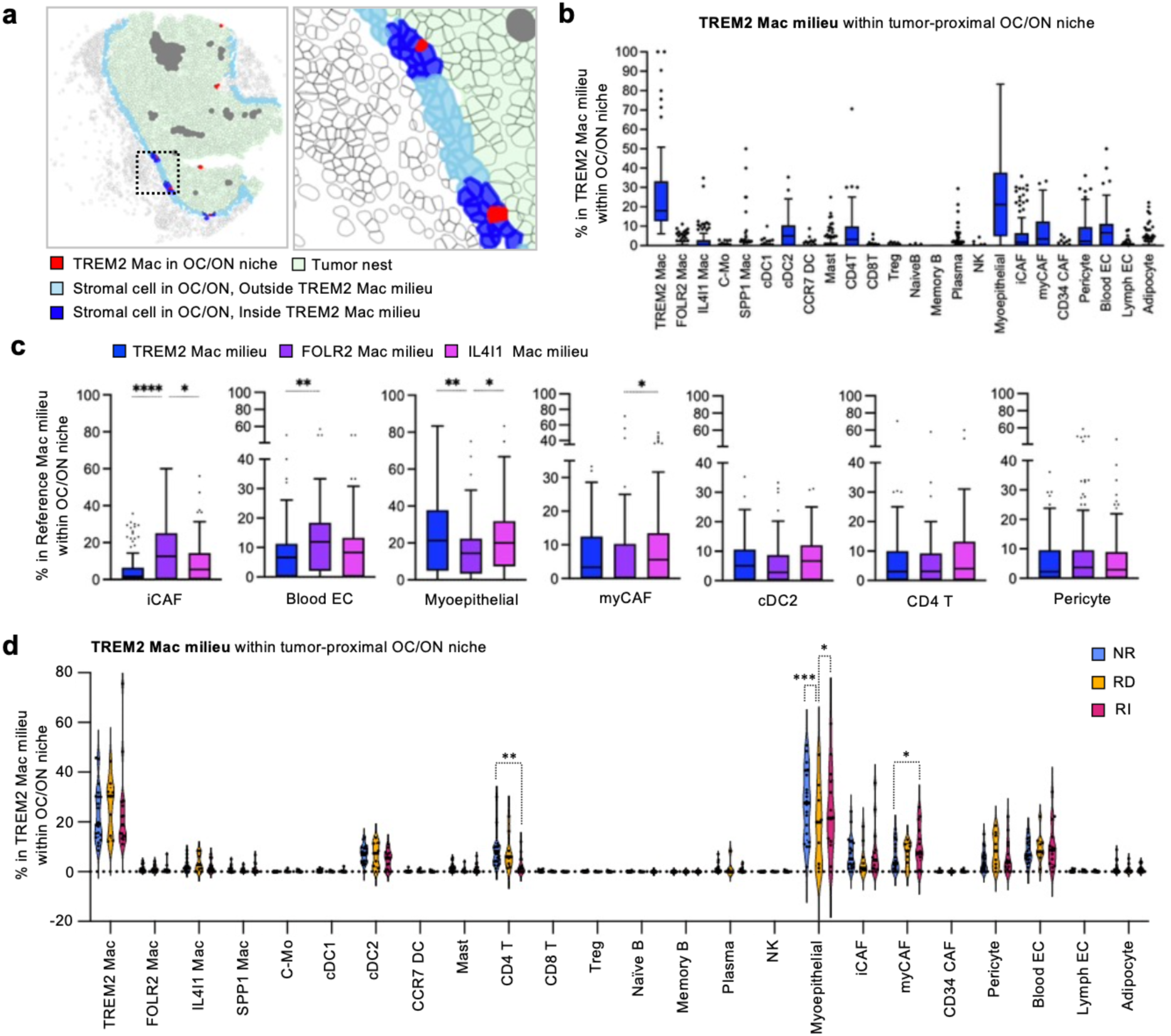
Stromal cell subsets neighboring TREM2^+^ macrophages within tumor-proximal niches. **a.** Representative image of the core shown in Fig. 3a displaying stromal cells within the TREM2 Mac milieu, defined as the area within four cell diameters of TREM2^+^ Macs, in the tumor-proximal OC/ON niches. High-magnification image of the selected area is shown on the right. **b.** Tukey box plots showing the proportions of annotated stromal cells within the TREM2 Mac milieu per core. Boxes represent the 25th–75th percentiles, with the median shown as a horizontal line. Whiskers extend to the most extreme data points within 1.5 × the interquartile range of the respective quartile. Values beyond the whiskers are plotted individually as outliers. **c.** Tukey box plots showing the proportions of annotated stromal cells within the milieus of TREM2^+^ Macs, FOLR2^+^ Macs, and IL4I1^+^ Macs in the OC/ON niches per core. Statistical differences among milieus were assessed using Kruskal–Wallis tests followed by Dunn’s multiple-comparison test (\**p* < 0.05, \*\**p* < 0.01, and \*\*\*\**p* < 0.0001). **d.** Violin plots showing the proportions of stromal cells within the TREM2 Mac milieu in the OC/ON niches across patient groups (NR, RD, and RI). Each dot represents one case. Statistical differences among patient groups were assessed by two-way ANOVA followed by Tukey’s multiple-comparisons test (\**p* < 0.05, \*\**p* < 0.01, \*\*\**p* < 0.001). Mac, macrophage; C-Mo, classical monocyte; DC, dendritic cell; Treg, regulatory T cell; NK, natural killer cell; CAF, cancer-associated fibroblast; EC, endothelial cell; OC, outside-nest/contact with epithelium; ON, outside-nest/near epithelium; NR, no recurrence; RD, recurrent DCIS; RI, recurrent IBC.

To determine whether this pattern was specific to the TREM2^+^ macrophage milieu, similar analyses were performed for the other major macrophage subpopulations within the tumor-proximal OC/ON niches, specifically FOLR2^+^ and IL4I1^+^ macrophages (**Supplementary Fig. 6a**), despite their lower abundance in these niches (**Fig. 3c**). The composition of the FOLR2^+^ macrophage milieu differed from that of the TREM2^+^ macrophage milieu (**Supplementary Fig. 6a, left**), with higher proportions of iCAFs and blood ECs and a lower proportion of myoepithelial cells (**Fig. 4c**). The proportions of myCAFs, cDC2, CD4^+^ T cells, and pericytes were comparable between the FOLR2^+^ and TREM2^+^ macrophage milieus. The IL4I1^+^ macrophage milieu more closely resembled the TREM2^+^ macrophage milieu than FOLR2^+^ macrophage milieu (**Supplementary Fig. 6a, right**). Similar to the TREM2^+^ macrophage milieu, the IL4I1^+^ macrophage milieu contained lower proportions of iCAFs and blood ECs and a higher proportion of myoepithelial than the FOLR2^+^ macrophage milieu. In addition, the proportion of myCAFs was higher in the IL4I1^+^ macrophage milieu than in the FOLR2^+^ macrophage milieu (**Fig. 4c**). These findings indicate that that the cellular composition of macrophage milieus varies according to macrophage subset and is not simply a reflection of the broader stromal cell distribution within the tumor-proximal niche.

Comparison of TREM2^+^ macrophage milieu composition across patient groups revealed that the proportion of myCAFs was significantly higher in RI than in NR cases, whereas CD4^+^ T cells were significantly lower in RI than in NR cases (**Fig. 4d**). Although the proportion of myoepithelial cells was lower in RD than in either NR or RI, no significant difference was observed between NR and RI. The proportions of other cell types, including TREM2^+^ macrophages, blood ECs, and cDC2, were comparable across patient groups. In contrast, the proportion of myCAFs within the FOLR2^+^ macrophage milieu was significantly lower in RI than in NR cases. In the IL4I1^+^ macrophage milieu, the proportion of myCAF was comparable across patient groups, whereas the proportion of iCAFs was higher in RI than in NR cases (**Supplementary Fig. 6b**).

Together, these findings indicate that invasive recurrence is associated with tumor-proximal enrichment of TREM2^+^ macrophages and with distinct changes in the cellular composition of their local milieus, characterized by increased myCAF and reduced CD4^+^ T cell representation.

### TREM2^+^ macrophage-epithelium crosstalk correlates with DCIS recurrence as IBC

We then explored cell-surface and secreted molecules that could mediate signaling from TREM2^+^ macrophages to neighboring epithelial cells, myCAFs, and CD4^+^ T cells. First, the snRNA-seq data on the DCIS cases shown in Figure 1 were reanalyzed using CellChat ^[20]^, with TREM2⁺ macrophages defined as ligand sources and epithelial cells, myCAFs, or CD4⁺ T cells as target populations expressing cognate receptors (**Fig. 5a**). Between TREM2^+^ macrophages and tumor epithelial (T-Epi) or normal epithelial (N-Epi) cells, LGALS9-CD44, SPP1-CD44, and GRN-SORT1 were the most probable ligand-receptor pairs (probability >0.01). The SEMA4C–PLXNB2 and CD99–PILRB pairs also exceeded a probability of 0.01 in N-Epi cells. Although integrin heterodimers, including ITGAV/ITGB1, ITGAV/ITGB5, and ITGA5/ITGB1 are also cognate receptors for SPP1, their interaction probabilities with SPP1 (<0.01) were lower than that of SPP1–CD44 in both T-Epi and N-Epi cells (0.05 and 0.04, respectively). In contrast, SPP1-ITGAV/ITGB1, SPP1-ITGAV/ITGB5, SPP1-ITGA5/ITGB1, and CD99-CD99 were the most probable ligand-receptor pairs between TREM2^+^ macrophages and myCAFs. LGALS6-CD44 had a probability of 0.01, lower than that observed in epithelial cells (0.04), whereas SPP1-CD44 and GRN-SORT1 had probabilities <0.01. Similar to the interaction between TREM2^+^ macrophages and epithelial cells, LGALS6-CD44, SPP1-CD44, and CD99-PILRB were the most probable ligand-receptor pairs between TREM2^+^ macrophages and CD4+ T cells. Additionally, 13 ligand-receptor pairs, including LGALS9-PTPRC, showed probabilities >0.01 in this cell-cell interaction (**Fig. 5a**). This analysis highlighted LGALS9, SPP1, GRN, and CD99 as candidate ligands expressed by TREM2⁺ macrophages that could mediate signaling to neighboring epithelial cells, myCAFs, or CD4⁺ T cells.

**Fig. 5.**
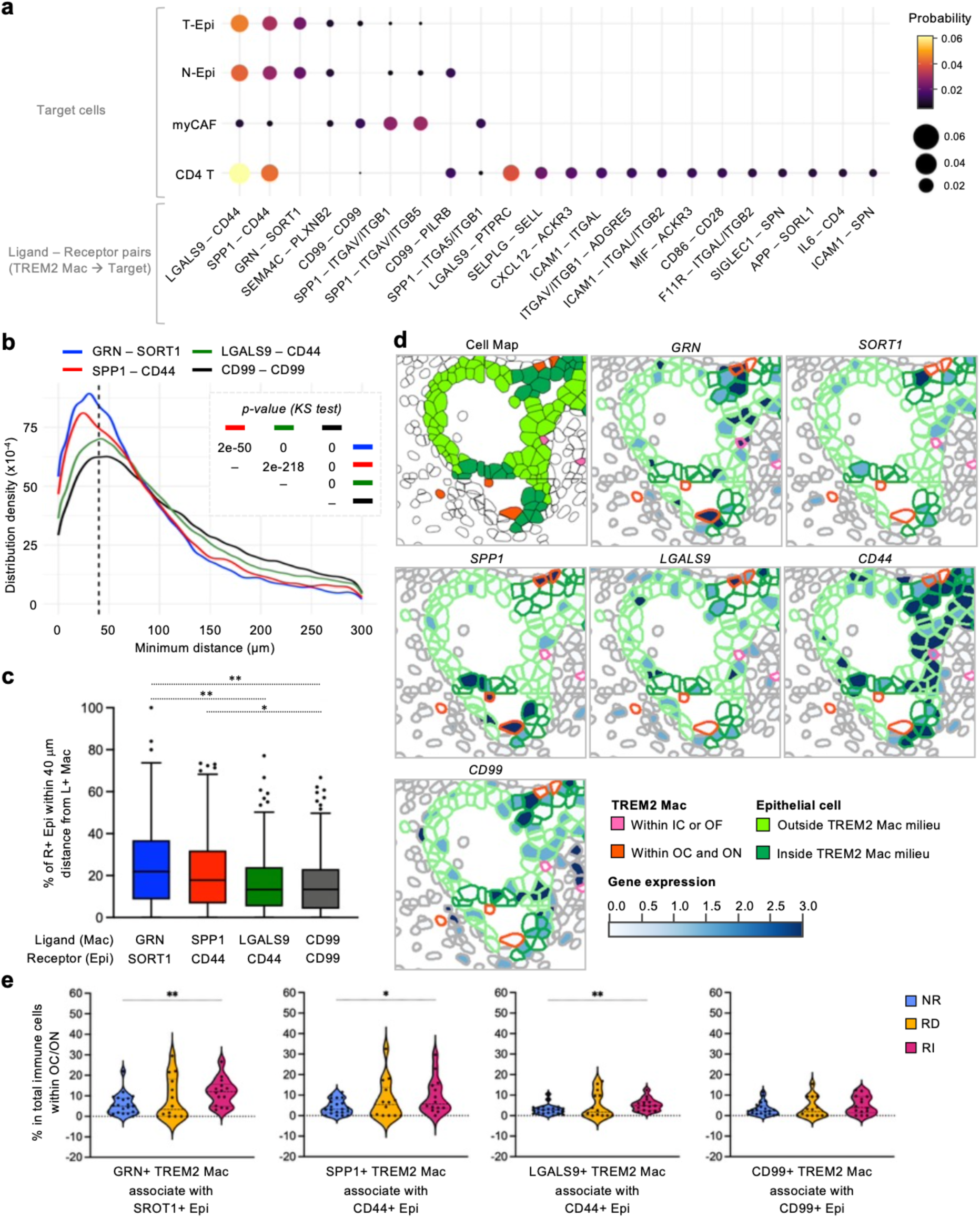
Potential interactions between epithelial cells and TREM2⁺ macrophages within the tumor-proximal niches. **a.** Bubble plot showing the probabilities of inferred ligand-receptor interactions between source (TREM2^+^ Macs) and target (indicated cells). Bubble size and color represent the interaction probabilities estimated from the snRNA-seq DCIS data (Fig. 1). **b.** Density plots showing the distribution of minimum distances between TREM2^+^ Macs and epithelial cells for the indicated ligand– receptor pairs. The x-axis represents the minimum cell-to-cell distance (µm), and the y-axis represents the estimated distribution density. The dashed vertical line indicates the 40-µm proximity threshold, corresponding to approximately four cell diameters. Statistical differences were assessed using the Kolmogorov-Smirnov (KS) test, with *p*-values shown in the insert table. **c.** Tukey box plots showing the proportions of indicated receptor-positive epithelial cells located within 40-µm of corresponding ligand-positive TREM2^+^ Macs per core. Boxes represent the 25th–75th percentiles, with the median shown as a horizontal line. Whiskers extend to the most extreme data points within 1.5 × the interquartile range of the respective quartile. Values beyond the whiskers are plotted individually as outliers. Statistical differences were assessed using Kruskal–Wallis tests followed by Dunn’s multiple-comparison test (\**p* < 0.05, \*\**p* < 0.01). **d.** Representative images showing the spatial distribution of tumor nest epithelial cells and surrounding TREM2^+^ Macs (Cell Map), together with the expression of the indicated genes. TREM2^+^ Macs within the tumor-proximal (OC/ON) and tumor-distal (OF) niches are shown in red and pink, respectively. Epithelial cells within and outside the TREM2^+^ Mac milieu are shown by dark and light green, respectively. Gene expression levels are indicated by a white-to-blue gradient. **e.** Violin plots showing the proportion of ligand-positive TREM2⁺ Macs spatially associated with corresponding receptor-positive epithelial cells within 40 µm, among all immune cells within the OC/ON niche, across patient groups (NR, RD, and RI). Each dot represents one case. Statistical differences were assessed using Mann–Whitney test (*p < 0.05, **p < 0.01). GRN, granulin; SORT1, sortilin-1; SPP1, secreted phosphoprotein-1; LGALS9, galectin-9; OC, outside-nest/contact with epithelium; ON, outside-nest/near epithelium; OF, outside-nest/far from epithelium; NR, no-recurrence; RD, recurrent DCIS; RI, recurrent IBC.

To assess whether these interactions were spatially plausible, we next examined the proximity between TREM2⁺ macrophages expressing these ligands and cells expressing the corresponding receptors using the DCIS spatial transcriptomics data. Specifically, we identified receptor-positive target cells across cores and quantified the minimum spatial distance to the nearest TREM2⁺ macrophage expressing the corresponding ligand. For interactions between TREM2^+^ macrophages and epithelial cells, the mode of the distance distribution was within 40 μm for all ligand–receptor pairs, with significantly shorter distances observed for GRN–SORT1 and SPP1–CD44 than for the other pairs (**Fig. 5b**). The same analyses performed separately for T-Epi and N-Epi cells revealed essentially the same distribution patterns as those observed for total epithelial cells (data not shown). For each core, we further calculated the proportion of receptor-positive epithelial cells within 40 µm of ligand-positive TREM2^+^ macrophages (**Fig. 5c**). GRN–SORT1 showed the highest proportion, with a median of 22% of SORT1-positive epithelial cells meeting this criterion, followed by SPP1– CD44 (18%), LGALS9–CD44 (13%), and CD99–CD99 (13%). For interactions between TREM2^+^ macrophages and myCAF, similar distribution trends were observed, although the modes of the distance distributions were greater than 40 μm and the median proportions of receptor-positive myCAFs located within 40 µm of ligand-positive TREM2^+^ macrophages were lower for all ligand–receptor pairs than those observed for TREM2^+^ macrophage-epithelial interactions (**Supplementary Fig. 7a,b**). Of note, the distribution patterns for the other SPP1 receptors, ITGAV/ITGB1 and ITGAV/ITGB5, were identical to those observed for CD44 (data not shown). For interactions between TREM2^+^ macrophages and CD4+ T cells, the distribution patterns differed substantially from those observed for epithelial cells or myCAFs, with the mode of the distance distribution exceeding 40 μm for all ligand–receptor pairs (**Supplementary Fig. 7c**). These results suggest that several inferred ligand–receptor pairs, particularly GRN–SORT1, SPP1–CD44, and LGALS9–CD44, are spatially plausible between TREM2⁺ macrophages and epithelial cells. Consistent with these findings, TREM2^+^ macrophages expressing GRN, SPP1, and LGALS9 were spatially associated with epithelial cells expressing their cognate receptors within tumor-proximal niche (**Fig. 5d**).

We next investigated whether the abundance of TREM2⁺ macrophages involved in the spatially plausible ligand–receptor interaction was associated with DCIS recurrence status. To this end, we quantified ligand-positive TREM2^+^ macrophages within the tumor-proximal OC/ON niches that were spatially associated with receptor-positive epithelial cells (i.e., within 40 μm) and compared their proportions among total immune cells across patient groups (**Fig. 5e**). The proportion of GRN-positive TREM2⁺ macrophages associated with SORT1-positive epithelial cells and proportion of SPP1-positive TREM2⁺ macrophages associated with CD44-positive epithelial cells were significantly higher in RI than in NR cases (**Fig. 5e**), with levels comparable to the proportion of total TREM2⁺ macrophages within the tumor-proximal niche (**Fig. 3d**). The proportion of LGALS9-positive TREM2⁺ macrophages associated with CD44-positive Epi cells was also higher in RI cases, although its abundance was lower than total TREM2⁺ macrophages. In contrast, no significant difference was observed in the proportion of CD99-positive TREM2⁺ macrophages associated with CD99-positive epithelial cells.

Together, these findings identify GRN–SORT1, SPP1–CD44, and LGALS9–CD44 as the most plausible candidate ligand–receptor interactions between TREM2⁺ macrophages and epithelial cells within the tumor-proximal niche. Notably, the abundance of spatially associated ligand-positive TREM2⁺ macrophages strongly correlated with the overall abundance of TREM2⁺ macrophages (**Supplementary Fig. 7d**), suggesting that their increased abundance in RI cases largely reflects expansion of the overall TREM2⁺ macrophage population rather than selective enrichment of these specific interactions.

### Protein-level spatial mapping of macrophage markers associated with DCIS recurrence

Finally, we performed multiplex fluorescence immunostaining on serial sections of the same 6 TMAs used for spatial transcriptomics to validate the spatial localization of macrophage markers (IBA1, TREM2, and FOLR2) at the protein level. The expression of pan-cytokeratin was also detected across sections to define the tumor nests (**Fig. 6a**). Following quality control and filtering using the same exclusion criteria for spatial transcriptomics, a total of 224 cores from 15 RI, 16 RD, and 21 NR cases were retained. For spatial mapping of macrophage marker proteins, the TME was categorized into three spatial niches according to their relationship to tumor nests (**Fig. 6b**). The intra-tumoral (IT) niche comprised stromal cells located within tumor nests, the tumor-proximal (TP) niche comprised stromal cells within the four-cell-distance peri-tumoral region, and the tumor-distal (TD) niche comprised stromal cells located distal to tumor nests. In this classification, IT corresponds to IC, TP to OC+ON, and TD to OF niches in the spatial transcriptomic analyses. Across the cores, the proportion of IBA1⁺TREM2⁺ macrophages among all IBA1⁺ macrophage was significantly higher in niches closer to tumor nests (**Fig. 6c,d**). In contrast, the proportion of IBA1⁺FOLR2⁺ macrophage was significantly increased in niches farther from tumor nests, consistent with the transcriptomic data. We then examined the association between the spatially distinct distributions of macrophage subsets and DCIS recurrence status, including non-recurrence (NR), recurrence with DCIS (RD), and recurrence with IBC (RI). Within the IT and TP niches, the proportion of IBA1⁺TREM2⁺ macrophage was significantly higher in RI cases than in NR cases, whereas no significant difference was observed in the TD niche (**Fig. 6e**). These niche-dependent, recurrence-associated changes were not observed for IBA1⁺FOLR2⁺ macrophages.

We further investigated the protein expression of candidate ligands implicated in spatially plausible interactions between TREM2⁺ macrophages and neighboring epithelial cells. Among all cells expressing progranulin (PGRN, encoded by *GRN*), osteopontin (OPN, encoded by *SPP1*), and galectin-9 (GAL9, encoded by *LGALS9*), macrophages constituted median fractions of 20%, 46%, and 44%, respectively (**Supplementary Fig. 8a**). Notably, OPN and GAL9 were significantly enriched in TREM2⁺ macrophages compared to FOLR2⁺ macrophages (**Supplementary Fig. 8b**). While the abundance of GAL9⁺ TREM2⁺ macrophages correlated with invasive recurrence (**Supplementary Fig. 8c**), this association mirrored that of the total TREM2⁺ population (**Supplementary Fig. 8d**), suggesting GAL9 expression is a constitutive feature of these recurrence-associated cells rather than an independent driver of risk stratification. Moreover, the abundance of GAL9⁺ TREM2⁺ macrophages was lower than that of total TREM2⁺ population (**Fig. 6e**). In contrast, the levels of PGRN⁺ and OPN⁺ TREM2⁺ macrophages did not differ significantly between recurrence groups (**Supplementary Fig. 8c,d**).

Collectively, these findings indicate that while the abundance of the TREM2⁺ macrophage niche serves as a robust prognostic biomarker, the specific expression levels of individual ligands (PGRN, OPN, GAL9) do not provide additional stratification value. However, the consistent expression of these molecules by recurrence-associated TREM2⁺ macrophages supports their role as key mechanistic mediators of macrophage–epithelial crosstalk. Thus, PGRN, OPN, and GAL9 remain compelling therapeutic targets for disrupting the stromal signaling networks that drive DCIS recurrence as IBC.

**Fig. 6.**
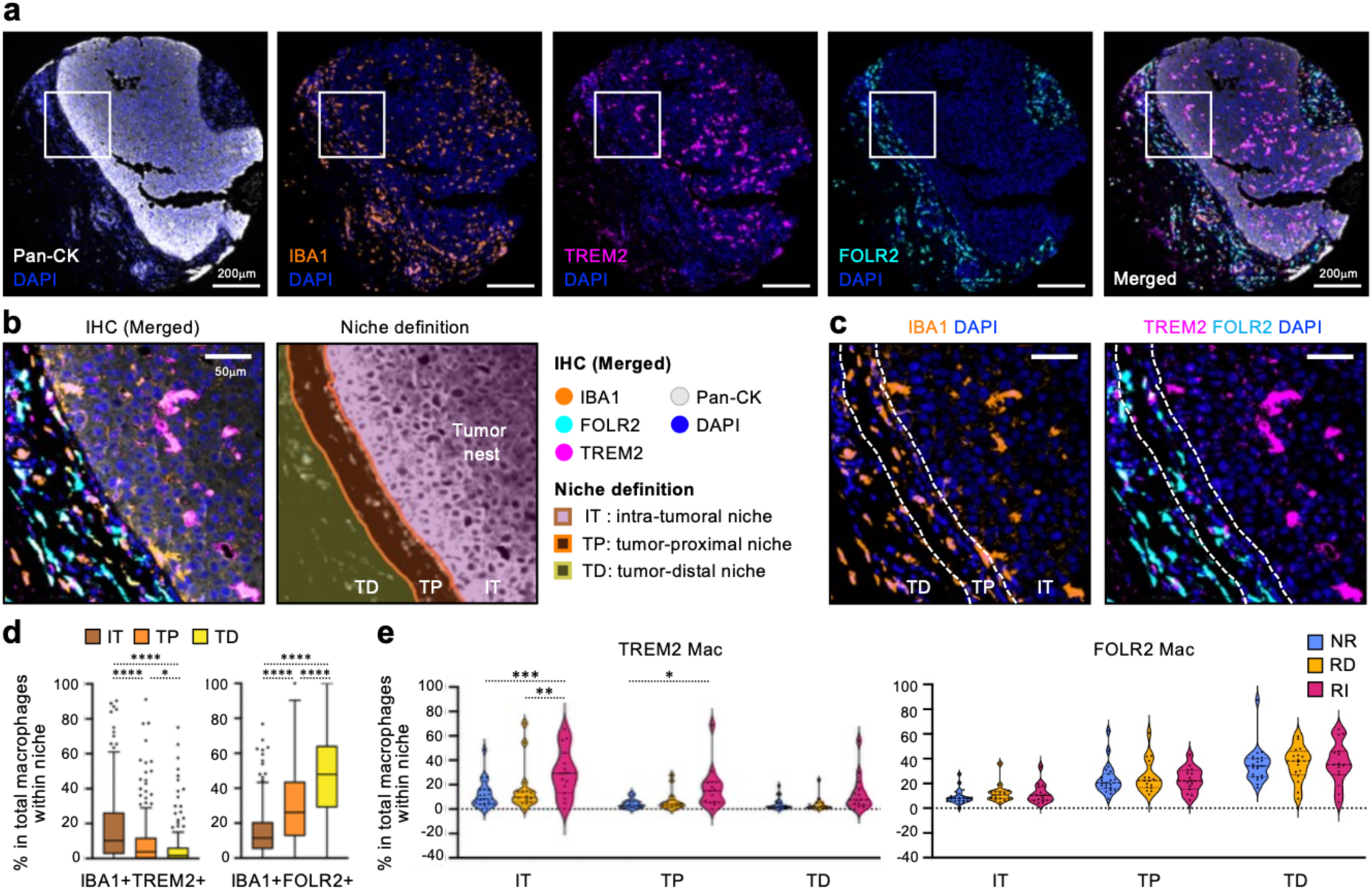
Spatial mapping of macrophage marker proteins within the DCIS tumor microenvironment. **a.** Representative multiplex immunostaining image of a core shown in Fig. 2e displaying the distribution of marker proteins for epithelial cells (Pan-CK) and macrophage subsets (IBA1, TREM2, and FOLR2). A merged image is also shown. **b.** High-magnification image of the selected area shown in a, displaying merged image (left) and defined niches relative to tumor nests(right). **c.** Localization of markers for pan-macrophage (IBA1) and macrophage subsets (TREM2 and FOLR2) within the area shown in b. **d.** Tukey box plots showing the proportions of macrophage subsets among all macrophages within each niche. Boxes represent the 25th–75th percentiles, with the median shown as a horizontal line. Whiskers extend to the most extreme data points within 1.5 × the interquartile range of the respective quartile. Values beyond the whiskers are plotted individually as outliers. Statistical differences across niches were assessed using Kruskal–Wallis tests followed by Dunn’s multiple-comparison test (\**p* < 0.05 and \*\*\*\**p* < 0.0001). **e.** Violin plot showing the proportion of macrophage subsets among all macrophages within each niche across patient groups (NR, RD, and RI). Each dot represents one case. Statistical differences among patient groups were assessed by two-way ANOVA followed by Tukey’s multiple-comparisons test (\**p* < 0.05, \*\**p* < 0.01, and \*\*\**p* < 0.001). CK, cytokeratin; IT, intra-tumoral niche; TP, tumor-proximal niche; TD, tumor-distal niche; NR, no recurrence; RD, recurrent DCIS; RI, recurrent IBC.

## Discussion

DCIS is a non-invasive breast tumor with diverse clinical outcomes, highlighting the need to identify biomarkers that stratify the risk of recurrence and progression to IBC, as well as therapeutic targets to prevent these events. Although previous studies have suggested that the TME may influence the clinical behavior of DCIS, its cellular composition and spatial organization remain understudied. In this study, we integrated snRNA-seq and spatial transcriptomics to characterize the cellular and spatial landscape of DCIS. We revealed a gradient-like TME in which macrophage and fibroblast populations occupied spatially defined niches relative to the tumor epithelium. Notably, increased abundance of TREM2^+^ macrophages within tumor-proximal niches was associated with subsequent recurrence as IBC. We further identified spatially plausible interactions between TREM2^+^ macrophages and neighboring epithelial cells, particularly the GRN–SORT1 and SPP1–CD44 ligand-receptor pairs, which may contribute to the modulation of epithelial behavior. These findings provide spatially resolved insights into the TME of DCIS and highlight TREM2^+^ macrophages as potential biomarkers for identifying patients at increased risk of progression to invasive disease and as potential targets for future therapeutic strategies (**Fig. 7**).

**Fig. 7.**
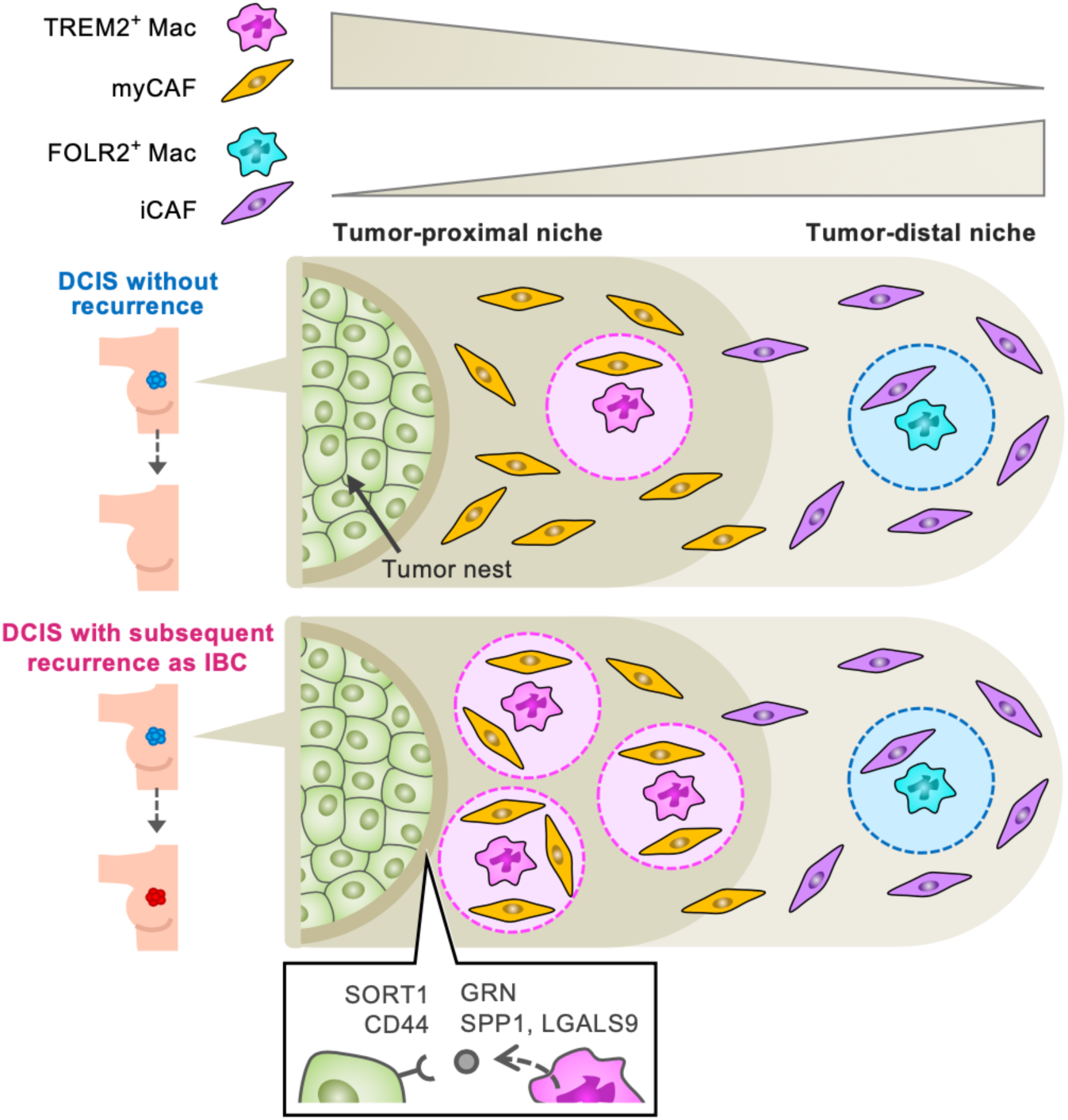
Graphical summary of this study. Spatial cell mapping revealed a gradient-like organization of the DCIS microenvironment. TREM2⁺ Macs and myCAFs were enriched in the tumor-proximal niche, whereas FOLR2⁺ Macs and iCAFs were preferentially localized in the tumor-distal niche. TREM2⁺ and FOLR2⁺ Mac subsets occupied distinct multicellular milieus characterized by a higher proportion of myCAFs and iCAFs, respectively. Notably, greater abundance of TREM2⁺ Macs in the tumor-proximal niche, as well as a higher proportion of myCAFs within the TREM2⁺ Mac milieu, was associated with recurrence as IBC. Ligand–receptor pair analysis further identified GRN–SORT1, SPP1–CD44, and LGALS9–CD44 as spatially plausible signaling interactions between TREM2⁺ Macs and epithelial cells. DCIS, ductal carcinoma *in situ*; IBC, invasive breast cancer; Mac, macrophage; CAF, cancer-associated fibroblast; TREM2, triggering receptor expressed on myeloid cells 2; FOLR2, folate receptor 2; myCAF, myofibroblastic CAF; iCAF, inflammatory CAF; GRN, progranulin; SORT1, sortilin 1; SPP1, secreted phosphoprotein 1; LGALS9, galectin 9.

A major finding of this study was the distinct spatial organization of TREM2^+^ and FOLR2^+^ macrophages within DCIS lesions. TREM2^+^ macrophages were preferentially localized within tumor nests and in tumor-proximal regions immediately surrounding epithelial structures, whereas FOLR2^+^ macrophages were rarely observed within tumor nests and progressively increased in abundance with increasing distance from tumor epithelium. Intriguingly, a similar spatial localization pattern has been reported in advanced breast cancer, where TREM2^+^ macrophages localize close to tumor nests, whereas FOLR2^+^ macrophages are largely located in the tumor stroma and rarely found within tumor nests ^[21]^. In this late-stage breast cancer study ^[21]^, TREM2^+^ and FOLR2^+^ macrophage subsets were distinguished by their differential expression of genes such as *TREM2*, *SPP1*, *CADM1*, *C3*, and *FN1* in TREM2^+^ subset, and *FOLR2*, *SEPP1*, *SLC40A1*, *MRC1*, *LYVE1*, and *CD163* in FOLR2^+^ subset. In our snRNA-seq dataset, including DCIS samples, differential gene expression analysis showed that 524 genes, including *TREM2*, *SPP1*, *CADM1*, and *C3,* were more highly expressed in TREM2^+^ macrophages than FOLR2^+^ macrophages (logFC >1.5, adjusted P <0.01), whereas 341 genes, including *SLC40A1*, *MRC1*, *LYVE1*, and *CD163*, were more highly expressed in FOLR2^+^ macrophages (**Supplementary Table 4**). Thus, the transcriptional profiles of TREM2^+^ and FOLR2^+^ macrophages in DCIS samples are consistent with those of the corresponding macrophage populations described in advanced breast cancer. Together with their similar spatial localization patterns, these data indicate that the gradient-like spatial distribution of TREM2^+^ and FOLR2^+^ macrophages is already apparent at the pre-invasive stage. Importantly, the abundance of tumor-proximal TREM2^+^ macrophages, but not that of FOLR2^+^ macrophages, in primary DCIS was associated with subsequent recurrence as IBC, suggesting that the spatial organization of TREM2^+^ macrophage population may have functional implications for disease progression. This observation is consistent with findings in breast cancer ^[21]^, where TREM2^+^ macrophages were poorly represented in healthy breast tissues but increased in cancer tissues, whereas FOLR2^+^ macrophages were found in both healthy and cancer tissues.

Interestingly, recent spatial transcriptomic analysis of normal human breast tissue has shown that TREM2^+^ macrophages occupy a specialized ductal niche, whereas FOLR2^+^ macrophages are predominantly localized to the stromal compartment ^[22]^. This observation raises the possibility that the spatial segregation of TREM2^+^ and FOLR2^+^ macrophages observed in DCIS may reflect a pre-existing organization of macrophage subset distributions within normal breast tissue. Our data suggests that a shift in the TREM2+ and FOLR2+ gradient with an increase in TREM2+ macrophages abundance may drive progression. Consistent with this, a spontaneous breast cancer mouse model demonstrates that macrophage heterogeneity is driven by pre-existing tissue territories ^[23]^. In normal mammary glands, macrophages are predominantly Folr2⁺Lyve1⁺Mrc1⁺ stromal subsets, with a smaller population of Hes1⁺Cadm1⁺ ductal subsets. In pre-tumoral tissues, the ductal niche additionally comprises an emerging Trem2⁺Spp1⁺ subset. Upon progression into malignant tumors, the macrophage compartment is predominantly comprised of these Trem2⁺ and Hes1⁺ ductal subsets, with Folr2⁺ stromal subsets becoming a minor population. Interestingly, pseudotime trajectory analysis infers that both stromal and ductal lineages originate from monocytes within the tissues, suggesting that macrophage states are defined by pre-existing niches, and neoplastic transformation selectively expands the Trem2⁺ subpopulation. It has been reported that TREM2+ macrophages originate from tumor-infiltrating monocytes in human breast cancer and relevant mouse models ^[24,25]^. Therefore, these observations collectively support a model in which TREM2^+^ macrophages may have an intrinsic propensity to occupy an epithelial- or ductal-associated niche, and neoplastic transformation may subsequently increase their abundance, possibly through the recruitment of monocytes to this specific region.

The fibroblast compartment in DCIS was also spatially structured. Namely, myCAFs were enriched in the tumor-proximal niches whereas iCAFs were preferentially localized to the tumor-distal niche, suggesting that the spatial distribution of myCAFs and iCAFs was broadly coordinated with that of TREM2^+^ and FOLR2^+^ macrophages, respectively. Although the abundance of myCAFs or iCAFs across the spatial niches did not differ between non-recurrent cases and cases that did recur, the proportion of myCAFs located within four-cell-diameters of individual TREM2^+^ macrophage, which we defined as a local cellular milieu, was associated with recurrence as IBC. These observations indicate the importance of considering cellular populations not only in terms of their overall abundance but also in relation to their spatial context and neighboring cell types. Notably, the cellular compositions of the macrophage-centered milieu differed according to macrophage identity. The TREM2^+^ macrophage and IL4I1^+^ macrophage milieus within the tumor-proximal niche contained higher proportions of myCAF and myoepithelial cells than the FOLR2^+^ macrophage milieu, whereas the FOLR2^+^ macrophage milieu contained higher proportions of iCAFs and blood endothelial cells than the milieus of the other macrophage subsets. These findings indicate that individual macrophage subsets appear to occupy distinct multicellular milieus. Although it remains unclear whether macrophages shape these milieus or are preferentially maintained within pre-existing environments, the identification of macrophage subset-specific milieus provides a framework for understanding macrophage function in the DCIS TME. In particular, the biological significance and underlying mechanisms of the TREM2^+^ macrophage–myCAF association warrant further investigation.

In contrast to the stromal compartment, we did not identify significant recurrence-associated differences in the cellular composition or global transcriptional states of normal or tumor epithelial cells, consistent with previous reports ^[3]^. However, our data indicated that epithelial cells were embedded within markedly distinct microenvironments, particularly with respect to tumor-proximal TREM2^+^ macrophage abundance and local myCAF representation. These findings suggest that recurrence-associated epithelial biology may be influenced not only by intrinsic epithelial states but also by interactions with the surrounding stromal and immune compartments. Given the close spatial association between TREM2^+^ macrophages and epithelial cells, we therefore investigated potential molecular interactions between these populations. Comprehensive analysis of our snRNA-seq and spatial transcriptomics data identified GRN–SORT1, SPP1–CD44, and LGALS9–CD44 as spatially plausible signaling pathways from TREM2^+^ Macs to epithelial cells at the tumor-proximal niche. Moreover, these interactions were more prominent in cases that recurred as IBC than in non-recurrent cases, largely reflecting the expansion of the overall tumor-proximal TREM2^+^ macrophage population. These findings raise the possibility that increased abundance of TREM2^+^ macrophages may influence epithelial behavior through specific ligand–receptor interactions, providing a potential mechanistic link between the macrophage-rich microenvironment and recurrence-associated progression. Although these ligands and receptors have rarely been studied in DCIS, several studies have implicated them in breast cancer progression. For example, proguranulin (GRN) has been shown to signal through sortilin 1 (SORT1) in breast cancer cells, and high tumor co-expression of GRN and SORT1 has been associated with poorer breast cancer-specific survival, providing clinical support for a role of this axis in aggressive breast cancer phenotypes ^[26,27]^. In addition, sortilin 1 (SORT1) is increased in invasive breast carcinomas and contributes to breast cancer cell adhesion, migration, and invasion ^[28]^. Moreover, single-cell RNA sequencing on breast cancer samples identified SPP1-expressing macrophages interacting with CD44-expressing malignant epithelial cells, with this interaction further supported by multiplex immunofluorescence ^[29]^. Moreover, osteopontin (SPP1) is increased in recurrent breast cancer and promotes recurrent tumor growth in experimental breast cancer mouse models ^[30,31]^. Although the roles of galectin 9 (LGALS9) in breast cancer is comparatively less established, a single-cell study has implicated LGALS9-CD44 and SPP1-CD44 signaling networks as potential interactions between macrophage subtypes and breast cancer cells ^[32]^. Collectively, the enrichment of these signaling interactions within tumor-proximal niches in DCIS may contribute to subsequent recurrence as invasive disease. Functional experiments will therefore provide an important opportunity to establish whether these candidate interactions directly promote disease progression and to define their potential as therapeutic targets.

Among the spatial features examined in this study, the most prominent recurrence-associated alteration was the accumulation of TREM2^+^ macrophages in proximity to the tumor epithelium. Multiplex fluorescence imaging further validated this observation at the protein-level. Specifically, IBA1^+^TREM2^+^ macrophage subsets were preferentially localized within niches closer to tumor nests, and their proportion among all macrophages was significantly higher in cases that subsequently recurred as IBC. The concordance between transcriptomic and protein-level analyses supports the value of incorporating spatially resolved immune profiling into the pathological assessment of DCIS and highlights TREM2 as a promising candidate biomarker for identifying patients with DCIS at increased risk of subsequent recurrence. Although a subset of TREM2⁺ macrophages expressed the candidate ligands PGRN, OPN, and GAL9 at the protein level, the abundance of the PGRN⁺ or OPN⁺ subsets was not significantly associated with recurrence status. While GAL9⁺ TREM2⁺ macrophage abundance correlated with recurrence, this trend simply mirrored the expansion of the total TREM2⁺ niche. Consequently, stratifying TREM2⁺ macrophages by these ligands offered no additional discriminatory value beyond the broader IBA1⁺TREM2⁺ population. This observation suggests that the overall abundance and spatial localization of TREM2⁺ macrophages robustly capture the recurrence-associated phenotype, rendering further stratification by individual ligand expression unnecessary. From a translational perspective, the use of a low-plex marker combination may offer practical advantages for clinical implementation. Low-plex multiplex assays involving two to three markers are generally more amenable to routine pathological assessment, whereas increasing multiplexity can introduce additional technical, analytical, and interpretive challenges ^[33,34]^. In this context, the ability of the IBA1⁺TREM2⁺ phenotype to capture the recurrence-associated macrophage population without additional ligand markers supports the feasibility of a relatively low-plex, spatially resolved approach for risk stratification of patients with DCIS.

In conclusion, this study presents a spatially resolved landscape of the microenvironment in DCIS and identifies the accumulation of TREM2⁺ macrophages in proximity to tumor epithelium as a prominent feature associated with subsequent recurrence as IBC. This study also highlights GRN–SORT1, SPP1–CD44, and LGALS9–CD44 as spatially plausible signaling pathways that may contribute to epithelial cell behavior. These findings provide a basis for future studies to validate TREM2⁺ macrophages as biomarkers for identifying patients with DCIS at increased risk of recurrence and invasive progression, and to determine whether targeting macrophage–epithelial interactions may offer therapeutic opportunities to prevent the invasive progression of DCIS.

## Methods

### Human sample collection

Formalin-fixed paraffin-embedded (FFPE) tissues and anonymized clinical data were obtained from the Tissue Biobank at the Breast Cancer Unit, Western General Hospital. Ethics were approved by the Lothian NRS BioResource Committee under Tissue Act Scotland, 2006. The provision and use of FFPE samples and associated data were approved by the Lothian NRS BioResource RTB (REC ref-20/ES/0061), under sample request numbers 2164 (Cohort 1: snRNA-seq) and 2119 (Cohort 2: TMA/spatial transcriptomics). Cohort 1, used for snRNA-seq, comprised primary tumors diagnosed as low-grade DCIS (n = 4), high-grade DCIS (n = 4), or stage I IBC (n = 4). An independent cohort 2, used for spatial transcriptomics and immunostaining, comprised primary DCIS tissues from patients diagnosed between 2000 and 2010 who remained recurrence-free until the end of follow-up (n = 22), or who subsequently developed recurrent DCIS (n = 16) or IBC (n = 15) following surgery. All samples were obtained from treatment-naïve patients with no history of chronic steroid use and contained no areas of comedo necrosis. Detailed information of these cohorts can be found in Supplementary Tables 1 and 2.

### Nuclei extraction

For snRNA-seq, nuclei were isolated from FFPE sections using the snPATHO-seq technique ^[35]^. For each sample, two 25-µm FFPE curls were collected in RNase-free tubes, deparaffinized in reagent grade xylene (Millipore, #214736) three times for 10 min, rehydrated through decreasing concentrations of ethanol (100%, 100%, 70%, 50%, and 30%; VWR, #83813.360DP), and washed in RPMI1640 media (Thermo Fisher Scientific, #11875093). The curls were physically disrupted using a pestle (Fisher Scientific, #12-141-364) in 100 µl of dissociation solution consisting of 1mg/ml Liberase (Roche, #5401119001), 1mg/ml Collagenase D (Roche, #11088858001), and 1U/µl RNase inhibitor (Thermo, #EO0382) in RPMI1640 media. An additional 900 µl of dissociation solution was then added, and the samples were incubated for 60 min at 37 °C in a thermomixer at 800 rpm. The dissociated tissues were washed in 400 µl lysis buffer consisting of 1x Nuclei Ez lysis buffer (Sigma-Aldrich, #NUC101), 2% Ultrapure BSA (Thermo, #AM2618), and 1 U/µl RNase inhibitor. After centrifugation at 850 xg for 5 min at 4 °C, the pellet was resuspended in 1 ml of lysis buffer and incubated on ice for 10 min, with intermittent pipetting to homogenize the sample. The suspension including nuclei was filtered through a 70-µm filter (pluriSelect, #43-10070-50) to remove undigested tissue and washed three times by centrifugation at 850 ×g for 5 min at 4 °C, followed by resuspension in 0.5× PBS containing 0.02% BSA. Finally, the extracted nuclei were filtered through a 40-µm filter (pluriSelect, #43-10040-50) before library preparation.

### Library preparation and sequencing

Nuclei isolated from FFPE sections were quantified using a Luna FX7 automated cell counter (Logos Biosystems), and 80,000 nuclei were used as input for each sample. Gene expression libraries were generated using the Chromium Fixed RNA Kit for Human Transcriptome (10x Genomics, #PN1000476) according to the manufacturer’s instructions (Chromium Fixed RNA Profiling User Guide; CG000527). Probe hybridization was performed for 20 h, after which samples labeled with unique probe barcodes were pooled and processed using the Chromium Fixed RNA Profiling workflow for gel bead-in-emulsion (GEM) generation, reverse transcription, and library construction. Library amplification was performed using 13 PCR cycles for library A and 12 PCR cycles for libraries B and C. Amplified libraries were purified using SPRIselect magnetic beads (Beckman Coulter, # B23317). Libraries were dual-indexed using standard Illumina-compatible dual indices (10x Genomics, Dual Index TS set, #PN-1000251), with unique index combinations assigned to each pooled library. Library quality and fragment size distributions were assessed using the High Sensitivity D1000 DNA ScreenTape assay (Agilent). Twelve individually barcoded samples were pooled into three sequencing libraries based on library quality and concentration. Libraries were sequenced using the Novaseq X Plus system (Illumina) using 25B flow-cell, paired-end 2x 150bp chemistry and standard Illumina indexing primers with a 5% PhiX spike-in.

### snRNA-seq data processing

Raw sequencing fastq files were processed using Cell Ranger v9.0.1 (10x Genomics), with reference transcriptome (GRCh38-2024-A) and probe set (10x Genomics, Chromium Human Transcriptome Probe Set v1.1.0). The preprocessing pipeline was run on the 10x Genomics Cloud. Further analyses were carried in R v4.5.1, using the Seurat v5.4.0 package ^[36]^. Cells with more than 600 detected features and less than 25% mitochondrial gene content were retained for downstream analysis. Data from each sample was then subjected to the following functions with default parameters: NormalizeData(), FindVariableFeatures(), ScaleData(), RunPCA(). Layers were then integrated using the IntegrateLayers function and the Harmony method ^[37]^. After integration the follwoing were run: RunPCA(), FindNeighbors(), FindClusters() and RunUMAP() on the first 30 principal components.

### snRNA-seq cell type annotation

Seurat clusters were annotated into large cell types–epithelial cell, myeloid cell, lymphocyte, fibroblast and pericyte, endothelial cell, and adipocyte– based on the expression of key marker genes, *CD68, CSF1R, CD3E, MS4A1, CD79A, NCAM1, CD4, CD8A, PECAM1, ADIPOQ, ACTA2, COL1A1* and *EPCAM*. For each non-epithelial cell type, the data were subsetted and reprocessed using the same workflow as for the full dataset, including normalization, identification of variable features, scaling, PCA, nearest-neighbor graph construction, UMAP embedding, and clustering at a resolution of 3.0. Each cluster was subsequently annotated to define fine-grained cell types and subsets based on the expression of key marker genes, as shown in Fig. 1 and Supplementary Fig. 1. For epithelial cells, ploidy status was assessed using CopyKAT v1.1.0 ^[16]^ with the following parameters: ngene.chr=5, win.size=25, KS.cut=0.1, genome=”hg20”, distance=”euclidian”. Cells classified as “Diploid” by CopyKAT were annotated as normal epithelial cells, whereas those classified as “Aneuploid” were annotated as tumor epithelial cells.

### snRNA-seq differential gene expression and pathway enrichment analysis

Cells of each cell type of interest were pseudo-bulked at the sample level using the AggregateExpression() function in Seurat. The resulting summed counts were then analysed using DESeq2 ^[38]^ v1.52.0 to determine differential gene expression. Pathway enrichment was run using ClusterProfiler ^[39]^ v4.20.0 function enricher(), using the MSigDB Hallmark ^[40]^ gene signatures v2024.1.

### snRNA-seq ligand–receptor interaction analysis

Potential cell–cell communication was analyzed using CellChat ^[20]^ version 2.1.1 with the human ligand–receptor interaction database. Following preprocessing to retain expressed genes and supported ligand–receptor interactions, communication probabilities were estimated between annotated cell populations TREM2^+^ Mac, FOLR2^+^ Mac, N-Epi, T-Epi, myoepithelial, myCAF, iCAF and CD4^+^ T cell. Analyses were focused specifically on interactions in which TREM2^+^ macrophages acted as sender. Candidate interactions were further prioritized using differential-expression information to retain interactions supported by the transcriptional state of the relevant cell populations. Interaction significance was assessed using CellChat-derived P values, followed by Benjamini–Hochberg correction for multiple testing. Interactions with false discovery rate < 0.05 were considered statistically significant. Significant interactions involving TREM2^+^ macrophages were ranked according to communication probability. High-confidence interactions were additionally prioritized based on their communication probability higher than 0.01.

### Tissue microarray (TMA) preparation

Based on H&E-stained sections, regions containing both tumor ducts and surrounding stroma were identified, and representative areas were selected for each DCIS sample from the above-mentioned cohort (53 cases in total). Six cores (1 mm in diameter) were collected from each FFPE block and arrayed into six separate TMA blocks. Each TMA block therefore contained one core from each of the 53 cases. From each TMA block, 5-µm sections were mounted on Leica Bond Plus slides (Leica, #S21.2113.A) and baked overnight at 60°C before processing for spatial transcriptomics.

### Spatial transcriptomic (ST) data acquisition

Spatial transcriptomic profiling was performed using the CosMx Spatial Molecular Imaging (SMI) platform according to the manufacturer’s instructions (Bruker/NanoString, *CosMx SMI Manual Slide Preparation for RNA Assays*, MAN-10184-04). TMA sections were processed using the default experimental conditions recommended for breast cancer tissue, including 15-min target retrieval, permeabilization with proteinase K (3 µg/mL) for 30 min at 40°C, and fiducial preparation at 0.001%. Gene expression profiling was performed using the CosMx Human 6K Discovery Panel (Bruker/NanoString, #121500041), and immunohistochemistry (IHC) for cell segmentation was performed using the CosMx Human Universal Cell Segmentation Kit for RNA (Bruker/NanoString, #121500020), comprising DAPI, CD298/B2M, Pan-CK, CD45, and CD68 markers. Image acquisition was performed on the CosMx SMI instrument across six slides, with field of view (FOV) measuring 0.51 × 0.51 mm, using pre-bleaching configuration C as recommended for breast cancer tissue.

### ST data processing and export

Raw CosMx SMI data were processed and decoded using the AtoMx Spatial Informatics Platform (v2.2.1) according to the manufacturer’s instructions (Bruker/NanoString, *CosMx SMI Data Analysis*, MAN-10162-06). The AtoMx analysis pipeline, including quality control, normalization, principal component analysis (PCA), cell typing, and cell type quality check, was performed using the default settings. Cells were segmented based on the nuclear and membrane markers included in the segmentation kit, and decoded probe counts were assigned to individual segmented cells. Cell types were annotated by supervised classification using a reference matrix generated from the DCIS snRNA-seq dataset, comprising 16 cell types. Cell mapping images and IHC images of all cores/FOVs, together with study statistics tables including the mean number of transcripts per cell for each FOV, were exported for quality check. The count matrix, cell metadata, and segmentation data (polygons) were also exported for downstream analyses using custom Python scripts.

### ST data quality control

Quality control was performed at both the FOV and cell levels. For FOV-level quality control, the mean number of transcripts per cell and core/FOV images were examined. FOVs were excluded from downstream analysis if they met any of the following criteria: (i) mean transcripts per cell < 200; (ii) absence of Pan-CK positive epithelial cells; (iii) tissue detachment.

Following this quality control and filtering, 175 cores comprising 689 FOVs were retained for downstream analysis. Cell-level quality control was subsequently performed using a custom Python script. The cell metadata, segmentation data, and expression count matrix exported from AtoMx were aligned using a unique cell identifier. Cells lacking corresponding expression or metadata records were excluded, and the correspondence between the expression matrix and cell metadata was verified before downstream analysis. For each cell, the number of detected RNA molecules (nCount) and detected genes/features (nFeature) was calculated as measures of RNA detection quality. The distributions of nCount and nFeature were assessed globally and across annotated cell populations, and their relationship was examined at the individual-cell level (Supplementary Fig. 2).

### ST niche analysis

Spatial relationships between stromal and epithelial cells were quantified using custom Python scripts. The cell metadata, segmentation data, and core/patient information were imported and integrated, and core-level information was merged with the cell metadata from FOVs designated for analysis. Cores containing fewer than 1% epithelial cells were excluded from subsequent spatial analyses. Cell segmentation polygons were reconstructed for individual cells from their global x/y polygon coordinates using Shapely and used for subsequent characterization of epithelial structures and stromal niches. First, epithelial cells were spatially grouped into contiguous epithelial structures to define tumor nests. Clusters containing at least six epithelial cells and having an area of at least six times the median epithelial-cell area were classified as tumor nests, whereas smaller epithelial structures were treated as fragments. Stromal cells outside tumor nests were then assigned to outer niches according to their polygon-based distance from the nearest tumor nest. Stromal cells located within 5 µm of a tumor nest, corresponding to approximately half the median stromal-cell diameter, were assigned to the outside nest/contact with epithelium (OC) niche. Cells located beyond 5 µm and within 40 µm distance, corresponding to approximately four cell diameters, were assigned to the outside nest/near epithelium (ON) niche, whereas cells beyond 40 µm were assigned to the outside nest/far from epithelium (OF) niche. Distances were calculated in pixels and converted to micrometers using a conversion factor of 0.12 µm per pixel. Next, tumor-nest epithelial polygons were united, with small gaps within nests filled while larger enclosed regions were preserved as holes. Holes (lumens) representing epithelial cell-free interior regions within tumor nests that remained after small-gap filling and had an area ≥ the median epithelial-cell area were defined as intra-tumoral ducts. Subsequently, stromal cells located within tumor nests or intra-tumoral ducts were assigned to the inside nest/contact with epithelium (IC) niche or the inside nest/duct (ID) niche, respectively, based on their centroid positions. Niche assignments were visually inspected at the tissue-core level to confirm consistency with the underlying segmentation and epithelial architecture (Fig. 3). The numbers of stromal cells within each niche, together with the numbers of normal- and tumor-epithelial cells within tumor nests, were quantified per core and subsequently integrated with patient and clinical-group metadata.

### ST neighborhood analysis

Stromal cells neighboring TREM2^+^ macrophages were quantified using custom Python scripts. Cells annotated as TREM2 Mac within the OC and ON niches were designated as reference cells. Stromal cells located within 40 µm distance of the nearest reference cell were assigned to the TREM2 macrophage milieu. Cell numbers and proportions within the milieu were calculated per core across TMA sections and subsequently integrated with patient and clinical-group metadata.

### ST ligand–receptor proximity analysis

Candidate ligand–receptor interactions were identified by CellChat using snRNA-seq data (described above), including SPP1–CD44, LGALS9–CD44, GRN–SORT1, and CD99–CD99. For each ligand–receptor pair, spatial proximity between ligand-positive TREM2^+^ macrophages (senders) and receptor-positive epithelial cells within tumor nests (receivers) was assessed in the CosMx spatial transcriptomic data using custom Python scripts. Cell metadata, segmentation data, expression count matrices, and tumor nest annotations were integrated using unique cell identifiers. Cells were considered positive for a given ligand or receptor when the corresponding transcript count was >0. For multi-subunit receptors, all required subunits had to have transcript counts >0. For each receptor-positive epithelial cell, the minimum geometric distance to the nearest ligand-positive TREM2^+^ macrophage within the same core was calculated from cell segmentation polygons and converted to micrometers using a conversion factor of 0.12 µm per pixel. The same spatial proximity analysis was performed for receptor-positive myCAFs and CD4^+^ T cells. Across six TMAs, spatial-distance distributions were assessed for each ligand–receptor pair and summarized by mean and median distance and by the proportion of receptor-positive cells located within 40 µm of a ligand-positive TREM2^+^ macrophage. In addition, ligand and receptor expression was visualized on a per-core basis together with cell-type identity and spatial niche localization. The number of ligand-positive TREM2^+^ macrophages located within 40 µm of receptor-positive cells was also quantified for each core and subsequently integrated with patient and clinical-group metadata.

### Multiplex immunohistochemistry (IHC)

Seven-plex fluorescence IHC was performed on FFPE sections using the Leica Bond Rx automated stainer. Serial sections (5 µm thick) were prepared from six TMA blocks used for ST analyses and baked for 18 h at 56 °C. Slides were deparaffinized twice in xylene for 5 min and rehydrated through graded ethanol (100%, 100%, 80%, 70%, 50%) followed by water, with each step performed for 20 s. Slides were subsequently subjected to sequential staining cycles using the Opal 6-plex detection system (Quanterix, #NEL871001KT). Each staining cycle consisted of antigen retrieval in BOND Epitope Retrieval Buffer 1 or 2 (ER1 or ER2; Leica, #AR0086 or #AR9640) for 20 min at 100°C, endogenous peroxidase blocking with 3% hydrogen peroxide for 10 min, incubation with a primary antibody for 60 min, incubation with a HRP-conjugated secondary antibody (Opal Polymer HRP Ms + Rb secondary antibody, Quanterix, #ARH1001A) for 30 min, and incubation with the corresponding Opal fluorophore (Opal 480, Opal 520, Opal 570, Opal 620, Opal 690, or Opal 780) diluted 1:150 in 1x Plus Amplification Diluent (Quanterix, #FP1609) for 10 min. Between each step, slides were washed three times for 5 min in 1x Tris-buffered saline (TBS, Corning, #46-012-CM) supplemented with 0.1% Tween-20 (Sigma, #P1379). In the final staining cycle for Opal 780 labeling, slides were incubated with Tyramide Signal Amplification-Digoxigenin (TSA-DIG, Quanterix, #NEL748) for 10 min following secondary antibody incubation. Slides were subsequently incubated in ER1 buffer for 20 min at 95 °C to strip the primary–secondary antibody complex while retaining the TSA-DIG signal, followed by incubation with an Opal 780-conjugated anti-digoxigenin antibody diluted 1:25 in antibody diluent/blocking buffer for 10 min. Following the final staining cycle, nuclei were counterstained with 1x Spectral DAPI (Quanterix, #FP1490) diluted in TBS, and slides were mounted using antifade mounting medium (Thermo, #P36961). Details of primary antibodies, including the corresponding antigen retrieval buffer and fluorophore, are provided in Supplementary Table 5.

### IHC data acquisition and analysis

Whole-slide multispectral imaging was performed using the Akoya PhenoImager HT (Quanterix, Vectra Polaris, v1.0.13). Subsequently, regions of interest (ROIs) were selected in Phenochart v1.1.0 (Quanterix) and exported to inForm v2.7.0 for spectral unmixing using the built-in algorithm. Unmixed images were stitched in Visiopharm v2023.09.4 and analyzed in QuPath v0.6.0 ^[41]^. Cells were segmented using InstanSeg ^[42]^ and phenotyped via threshold-based marker expression. Pan-cytokeratin-positive epithelial clusters were annotated as tumor nests. The tumor-proximal region was defined as the area extending 40 µm from the tumor nest, while the tumor-distal region was defined as the area beyond this proximal region. Adipose regions were excluded from analysis.

### Statistical analysis

Statistical analyses and data visualization were performed using GraphPad Prism v11.0.1 (GraphPad Software) or R v4.5.1. Unless otherwise specified, analyses were performed on a per-core basis, whereas comparisons among patient groups (NR, RD, and RI) were performed at the individual patient level, with integrated cores treated as the unit of analysis. Differences in cell-type proportions among multiple spatial niches were assessed using Kruskal–Wallis tests followed by Dunn’s multiple-comparisons tests. Differences in stromal cell proportions among patient groups were assessed using two-way analysis of variance (ANOVA) followed by Tukey’s multiple-comparisons test, whereas comparisons between two patient groups were performed using the Mann–Whitney U test. For spatial ligand–receptor analyses, distributions of minimum cell-to-cell distances between ligand- and receptor-positive cells were compared using the Kolmogorov–Smirnov (KS) test. Differences in the proportions of receptor-positive cells located within 40 μm of the corresponding ligand-positive cells were assessed using Kruskal–Wallis tests followed by Dunn’s multiple-comparisons tests. All statistical tests were two-sided, and *P* < 0.05 was considered statistically significant. Statistical significance is indicated by *, **, ***, and ****, corresponding to *P* < 0.05, *P* < 0.01, *P* < 0.001, and *P* < 0.0001, respectively.

## Data availability

Single-nucleus RNA sequencing and spatial transcriptomics data generated in this study will be made available upon publication.

## Code availability

All code developed and used in this study will be made available through a GitHub repository upon publication.

## Acknowledgements

This project was funded by Cancer Research UK (EDDPMA-Nov23/100058 to T.K), Breast Cancer Now (2024.11PR1774 to T.K), and Cancer Research Horizons (iTPA PIII159 to N.G). F.P. was supported by a Wellcome Trust Early Career Award (225021/Z/22/Z). N.G. was supported by a grant from Breast Cancer Now (2024.11PR1774). We gratefully acknowledge Vishad Patel and Craig Marshall (NHS Lothian, NRS Bioresource) for the generation and sectioning of TMA blocks; Meryam Beniazza and Viktoria Major (Single Cell & Spatial Biology Facility, Institute for Regeneration and Repair [IRR], University of Edinburgh [UoE]) for transcriptomic data acquisition; and Hazel Stewart, Melanie McMillan, and Justyna Cholewa-Waclaw (IRR Histology & Multiplexing Facility, UoE) for their support with multiplex immunostaining and image scanning.

## Author contributions

T.K., F.P., and N.G. conceived and designed the study. T.K. and F.P. supervised the study. C.M.P., A.T. and M.D. selected appropriate patient samples and provided clinical metadata. N.G. performed nuclei isolation for snRNA-seq. F.P. and N.G. analyzed the snRNA-seq data, and T.K., F.P., and N.G. analyzed the spatial transcriptomic data. N.G. performed multiplex immunostaining and image analysis. T.K., F.P., and N.G. wrote the manuscript. All authors reviewed and approved the final manuscript.

