## Supplementary Materials for "Deciphering the association of spatial landscapes of DCIS with recurrence as invasive breast cancer"

### Supplementary Figure 1

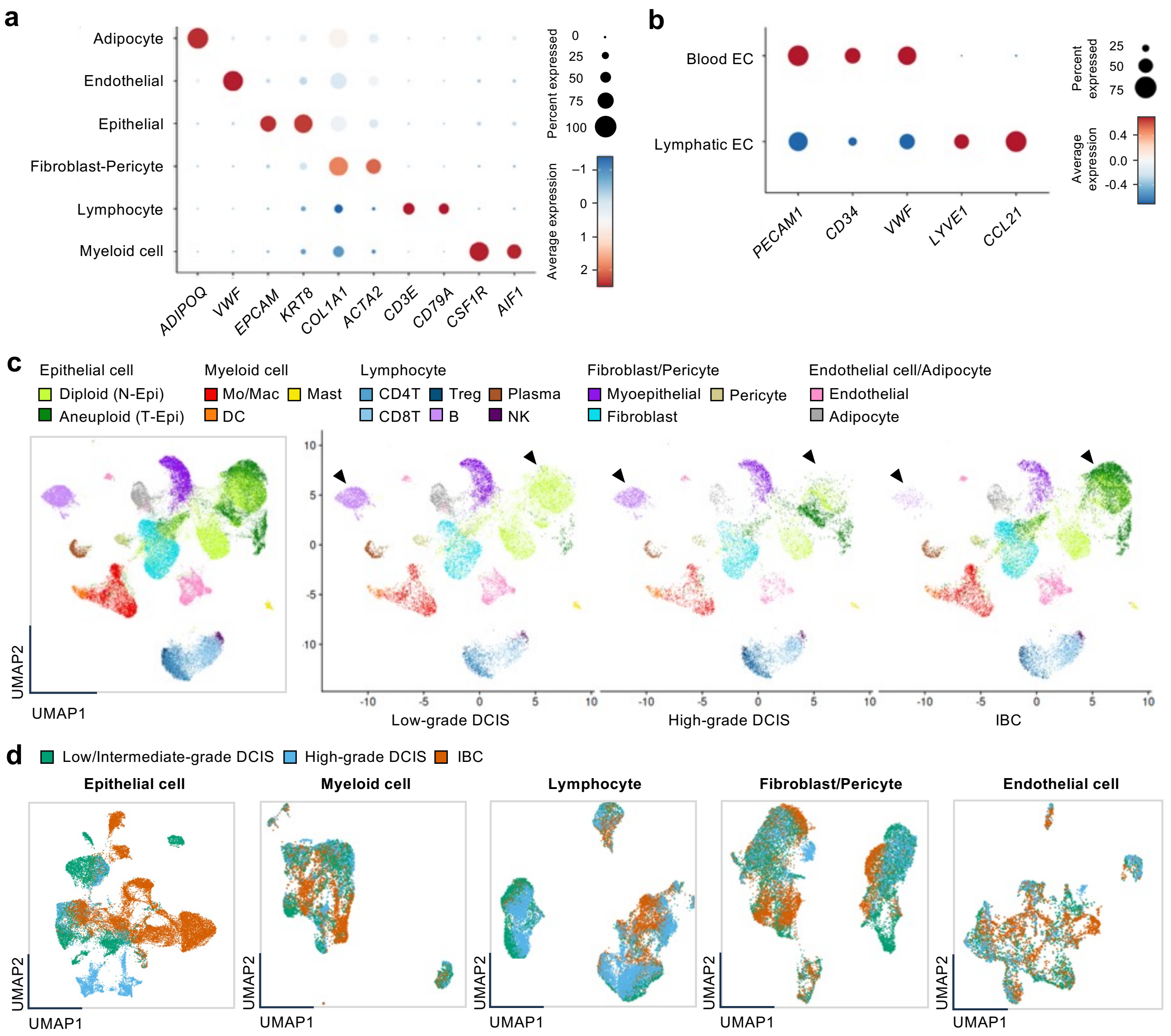


**Supplementary Fig. 1 Identification of cell types within DCIS. a.** The expression of lineage marker genes in major cell lineages shown in Figure 1b. Circle size and color indicate the percentage of cells expressing each gene and its average expression level, respectively. **b.** The expression of lineage marker genes in endothelial cell subpopulations shown in Figure 1d. **c.** UMAP plots showing subclusters identified by lineage-specific re-clustering (left) and their distribution in low/intermediate-grade DCIS, high-grade DCIS, and IBC (right). Arrow heads indicate clusters that were enriched in either DCIS or IBC. **d.** UMAP plots showing the distribution of epithelial and stromal subpopulations (presented in Figure 1c and d) across low-grade DCIS, high-grade DCIS, and IBC. Mac, macrophage; Mo, monocyte; DC, dendritic cell; Treg, regulatory T cell; NK, natural killer cell; DCIS, ductal carcinoma in situ; IBC, invasive breast cancer.

### Supplementary Figure 2

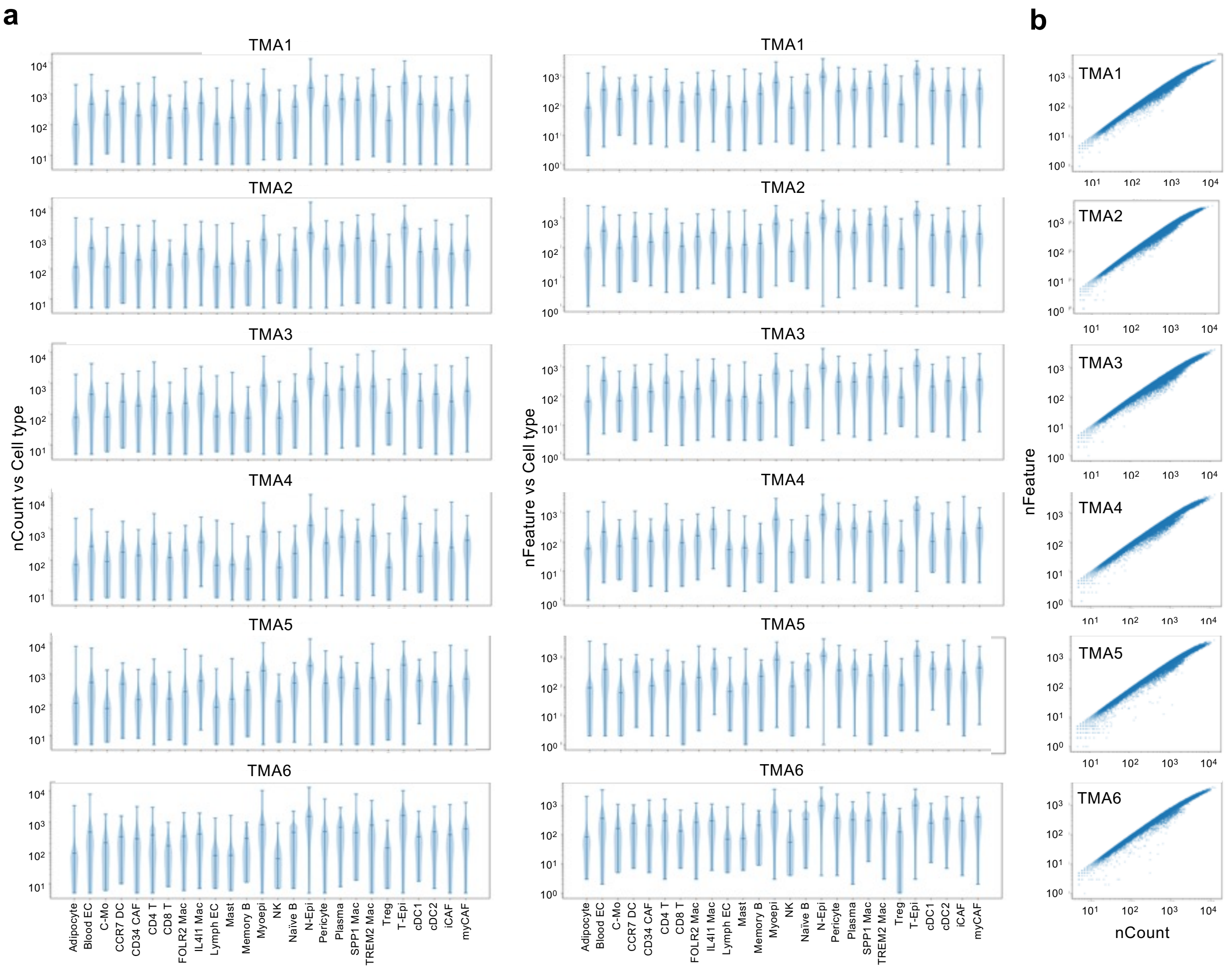


**Supplementary Fig. 2 Quality control assessment of cell-level RNA measurements. a.** Violin plots showing the distribution of total RNA counts (nCount; left) and the number of detected features (nFeature; right) across the annotated cell types. **b.** Scatter plots showing the relationship between nCount and nFeature for individual cells on a log–log scale. TMA, tissue microarray; N-Epi, normal epithelial cell; T-Epi, tumor epithelial cell; Mac, macrophage; C-Mo, classical monocyte; DC, dendritic cell; Treg, regulatory T cell; NK, natural killer cell; CAF, cancer-associated fibroblast; EC, endothelial cell; NR, no recurrence; RD, recurrent DCIS; RI, recurrent IBC.

### Supplementary Figure 3

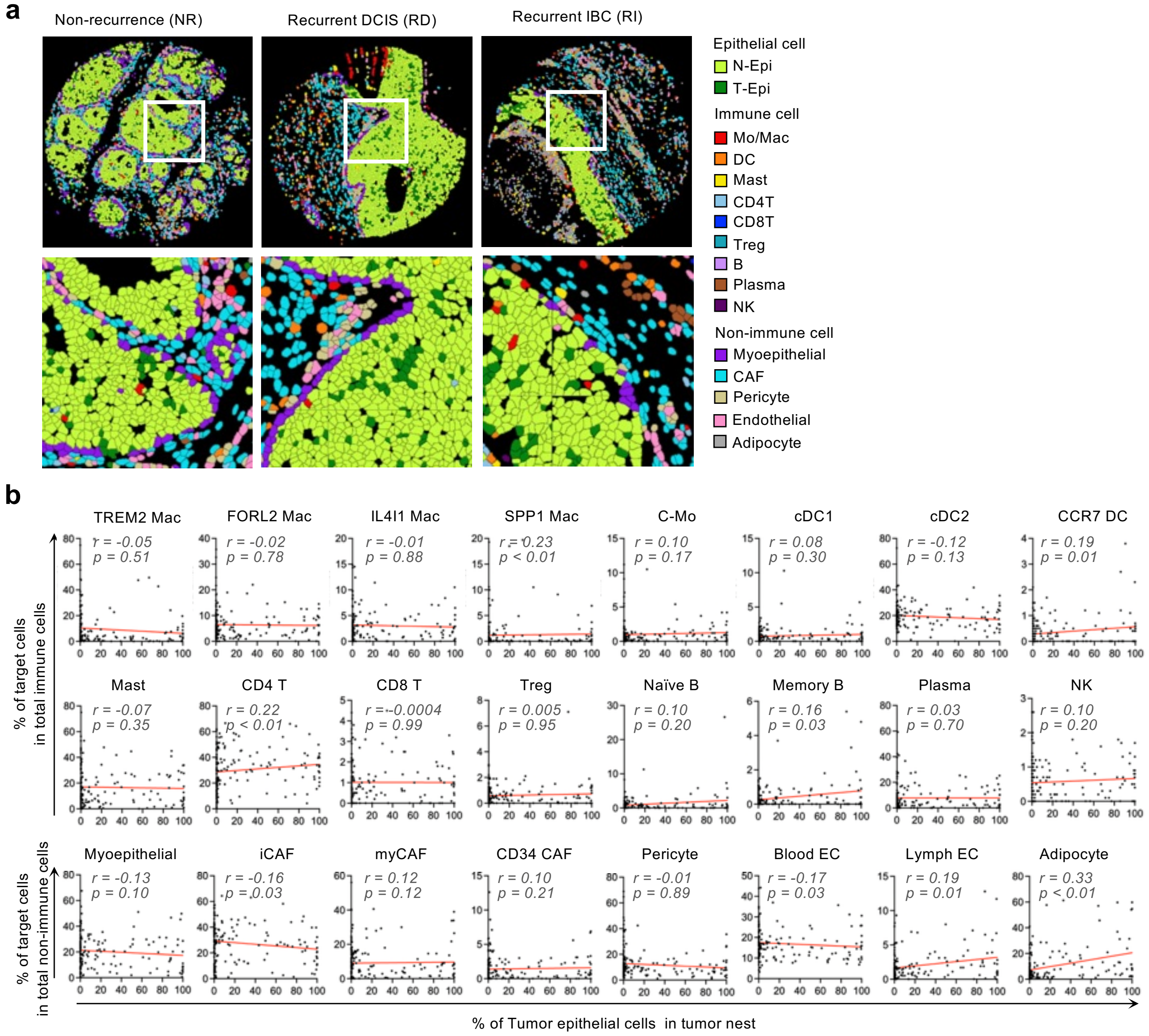


**Supplementary Fig. 3 Correlations between the abundance of tumor epithelial cells and stromal cell populations. a.** Representative spatial transcriptomic images of cores from each group (NR, RD, and RI) after annotation of major cell lineages. High-magnification images of selected areas are shown below each image. **b.** Scatter plots showing the relationships between the proportion of T-Epi cells within tumor nests and the proportions of defined stromal cell types per core. Spearman rank correlation coefficients (r) and p-values are shown in each plot. N-Epi, normal epithelial cell; T-Epi, tumor epithelial cell; Mac, macrophage; C-Mo, classical monocyte; DC, dendritic cell; Treg, regulatory T cell; NK, natural killer cell; CAF, cancer-associated fibroblast; EC, endothelial cell; NR, no recurrence; RD, recurrent DCIS; RI, recurrent IBC.

### Supplementary Figure 4

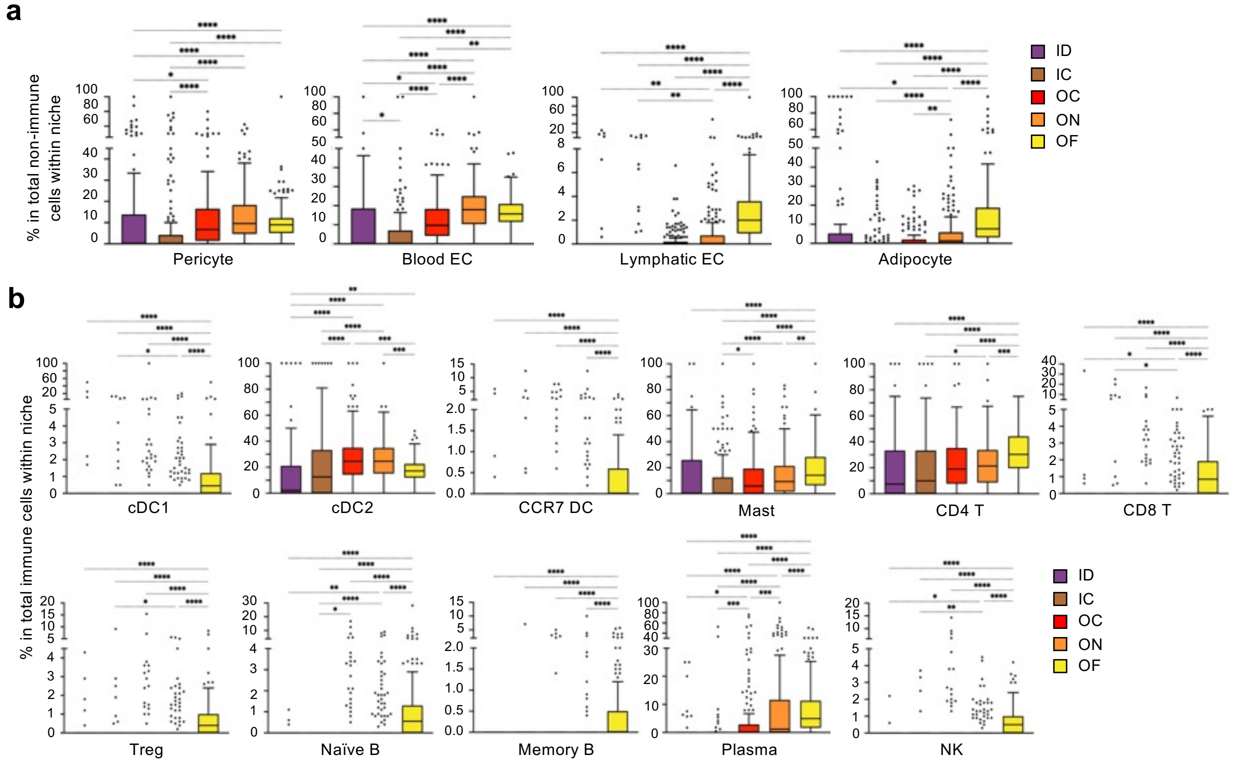


**Supplementary Fig. 4 Localization of stromal cell subsets in tumor-associated niches. a.** Tukey box plots showing the proportions of pericytes, endothelial cells, and adipocytes among non-immune stromal cells within each niche. **b.** Tukey box plots showing the proportions of immune cell subpopulations excluding monocytes/ macrophages among immune cells within each niche. Boxes represent the 25th–75th percentiles, with the median shown as a horizontal line. Whiskers extend to the most extreme data points within 1.5 × the interquartile range of the respective quartile. Values beyond the whiskers are plotted individually as outliers. Statistical differences across niches were assessed using Kruskal–Wallis tests followed by Dunn's multiple-comparison test (*$p < 0.05$, **$p < 0.01$, ***$p < 0.001$, and ****$p < 0.0001$). EC, endothelial cell; DC, dendritic cell; Treg, regulatory T cell; NK, natural killer cell; ID, inside-nest/duct; IC, inside-nest/contact with epithelium; OC, outside-nest/contact with epithelium; ON, outside-nest/near epithelium; OF, outside-nest/far from epithelium.

### Supplementary Figure 5

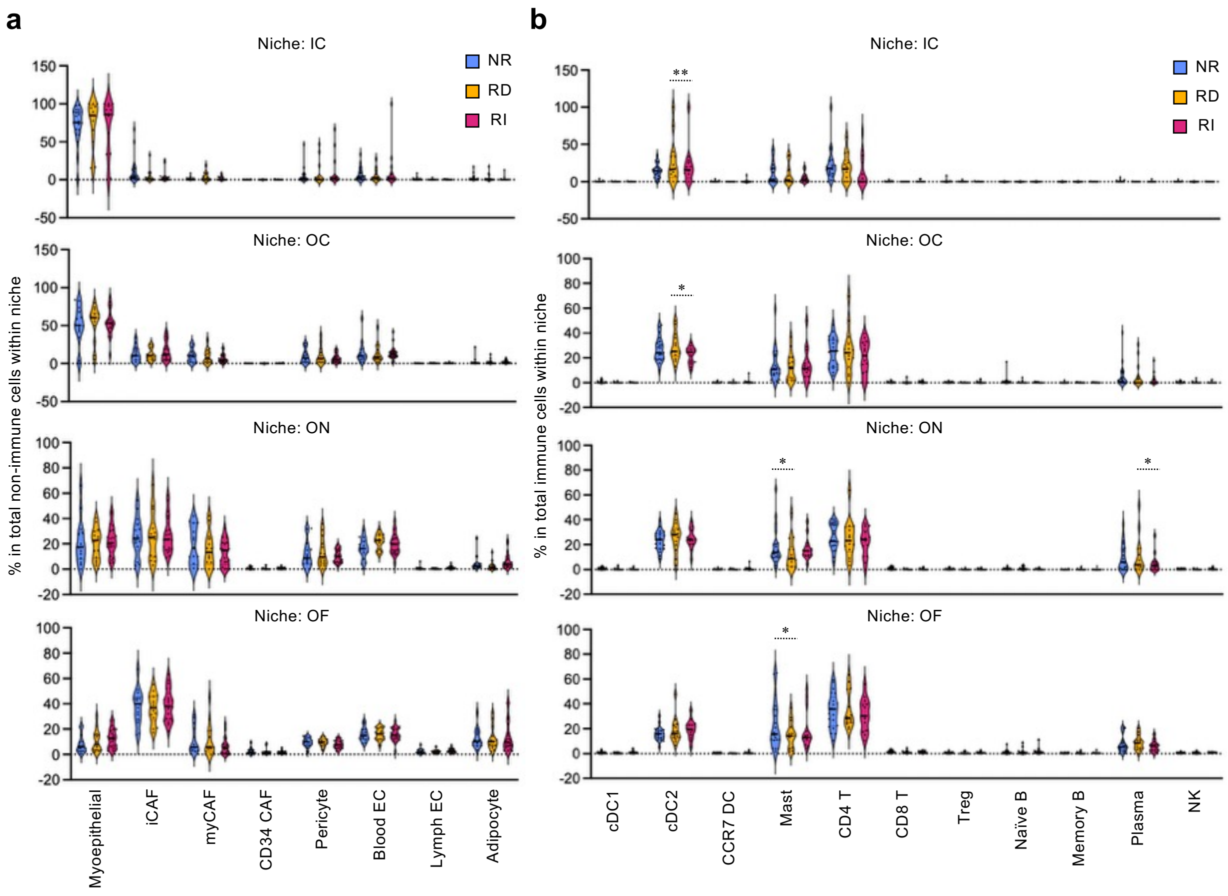


**Supplementary Fig. 5: Proportions of stromal cell populations across patient groups. a.** Violin plots showing he proportions of annotated cell types among non-immune cells within each niche across patient groups (NR, RD, and RI). Each dot represents one case. b. Violin plots showing he proportions of immune cell subpopulations excluding monocytes/ macrophages among immune cells within each niche across patient groups. Statistical differences among patient groups were assessed by two-way ANOVA followed by Tukey's multiple-comparisons test (*p < 0.05, **p < 0.01). CAF, cancer-associated fibroblast; EC, endothelial cell; DC, dendritic cell; Treg, regulatory T cell; NK, natural killer cell; ID, inside-nest/duct; IC, inside-nest/contact with epithelium; OC, outside-nest/contact with epithelium; ON, outside-nest/near epithelium; OF, outside-nest/far from epithelium; NR, no recurrence; RD, recurrent DCIS; RI, recurrent IBC.

#### Supplementary Figure 6

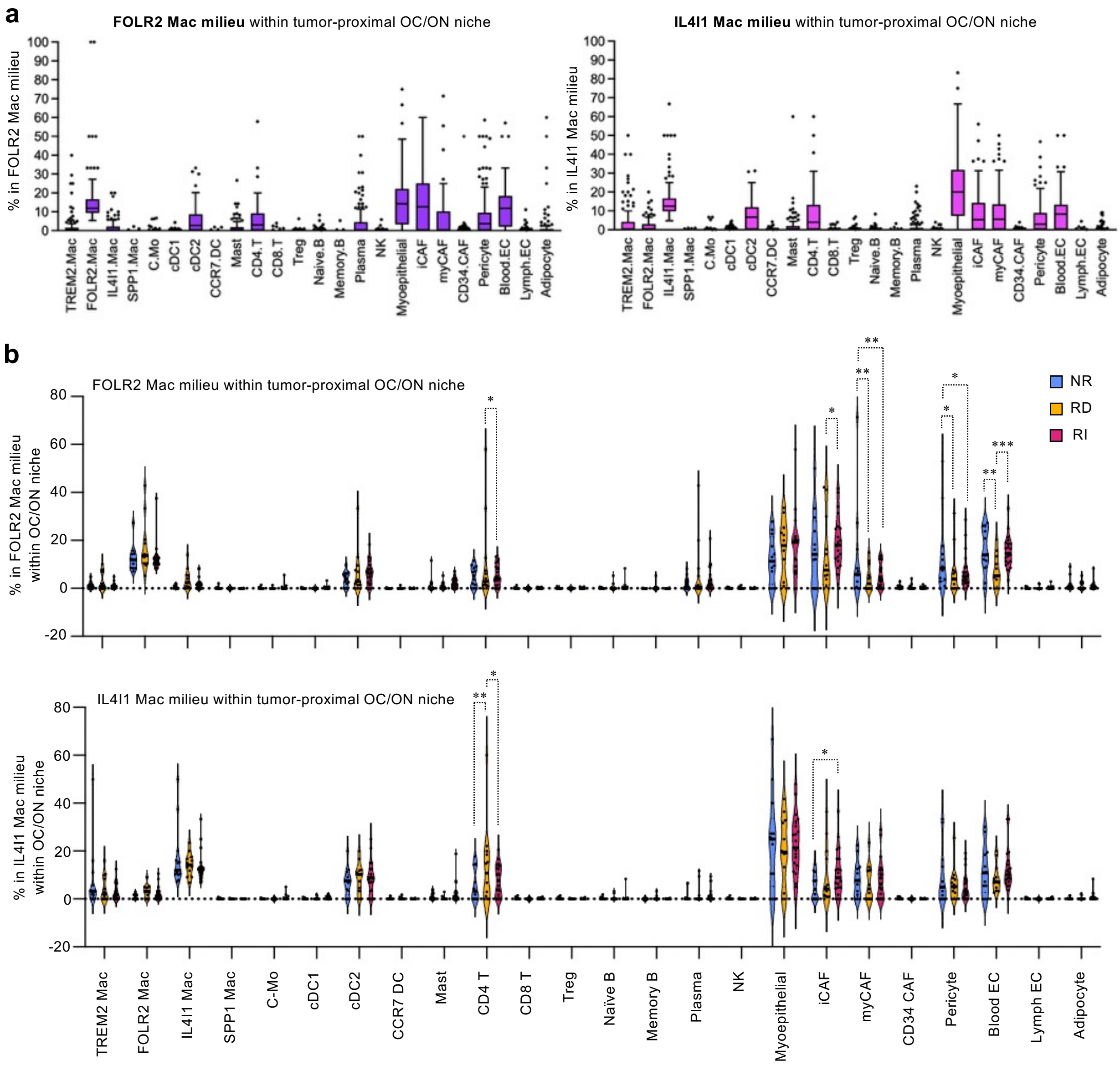


**Supplementary Fig. 6 Stromal cell subsets neighboring FOLR2+ macrophages and IL4I1+ macrophages within the tumor-proximal niches. a.** Tukey box plots showing the proportions of annotated stromal cells within the milieus of FOLR2+ Macs (left) and IL4I1+ Macs (right) per core. Boxes represent the 25th–75th percentiles, with the median shown as a horizontal line. Whiskers extend to the most extreme data points within 1.5 × the interquartile range of the respective quartile. Values beyond the whiskers are plotted individually as outliers. **b.** Violin plots showing the proportions of stromal cells within the FOLR2 Mac milieu (top) and IL4I1 Mac milieu (bottom) in the OC/ON niches across patient groups (NR, RD, and RI). Each dot represents one case. Statistical differences among patient groups were assessed by two-way ANOVA followed by Tukey's multiple-comparisons test (*$p < 0.05$, **$p < 0.01$, ***$p < 0.001$). Mac, macrophage; C-Mo, classical monocyte; DC, dendritic cell; Treg, regulatory T cell; NK, natural killer cell; CAF, cancer-associated fibroblast; EC, endothelial cell; OC, outside-nest/contact with epithelium; ON, outside-nest/near epithelium; NR, no recurrence; RD, recurrent DCIS; RI, recurrent IBC.

### Supplementary Figure 7

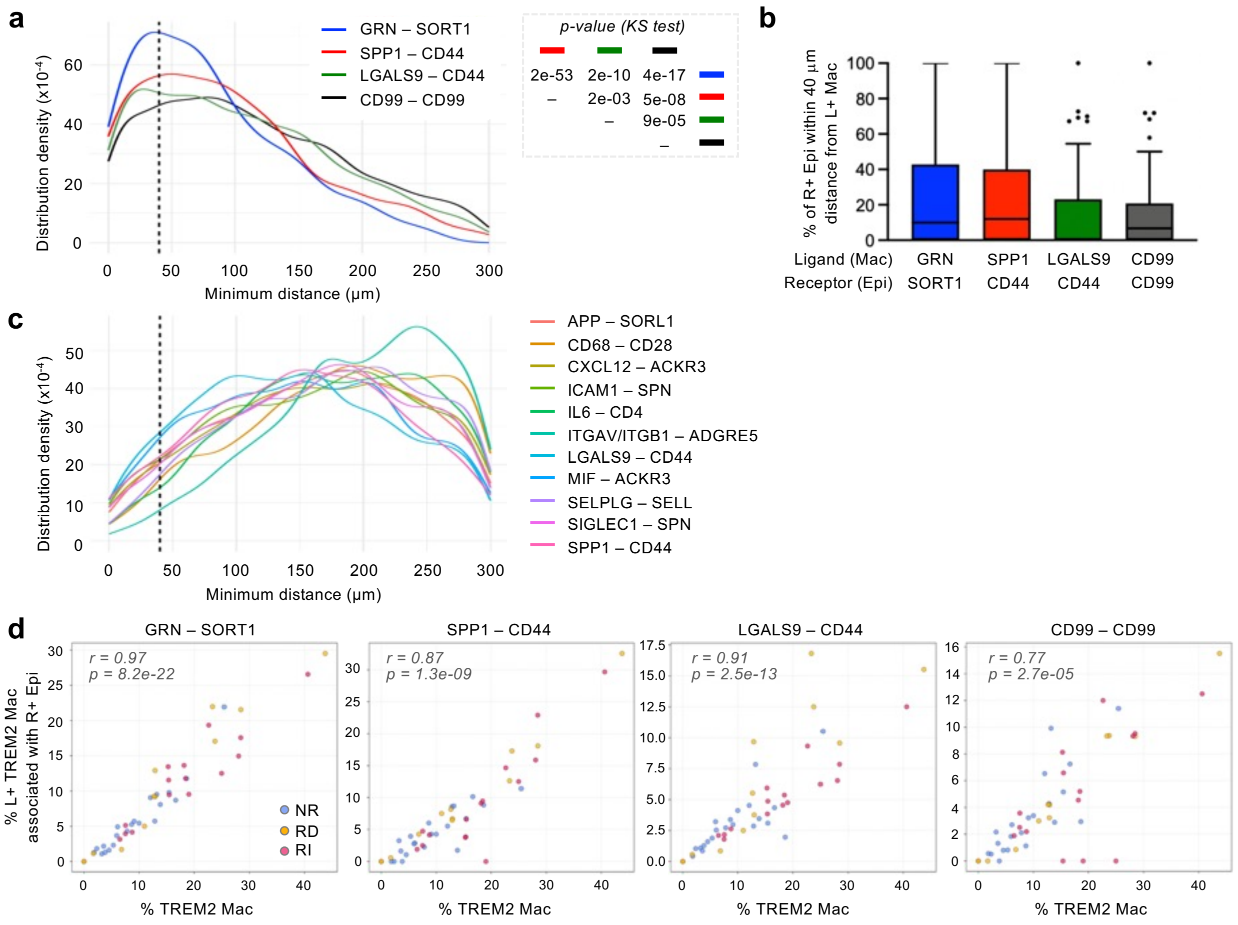


**Supplementary Fig. 7 Potential signaling interactions mediated by TREM2⁺ macrophages in the tumor-proximal niche. a.** Density plots showing the distribution of minimum distances between TREM2+ Macs and myCAFs for the indicated ligand–receptor pairs. The x-axis represents the minimum cell-to-cell distance (μm), and the y-axis represents the estimated distribution density. The dashed vertical line indicates the 40-μm proximity threshold, corresponding to approximately four cell diameters. Statistical differences among ligand-receptor pairs were assessed using the Kolmogorov-Smirnov (KS) test, with p-values shown in the insert table. **b.** Tukey box plots showing the proportions of indicated receptor-positive epithelial cells located within 40-μm of corresponding ligand-positive TREM2+ Macs per core. Boxes represent the 25th–75th percentiles, with the median shown as a horizontal line. Whiskers extend to the most extreme data points within 1.5 × the interquartile range of the respective quartile. Values beyond the whiskers are plotted individually as outliers. No statistical significances were detected by Kruskal–Wallis tests followed by Dunn's multiple-comparison test. **c.** Density plots showing the distribution of minimum distances between TREM2+ Macs and CD4+ T cells for the indicated ligand–receptor pairs. **d.** Correlations between the proportion of ligand-positive TREM2⁺ Macs spatially associated with corresponding receptor-positive epithelial cells within 40 μm (y-axis) and the proportion of TREM2+ Macs (x-axis), both calculated among all immune cells within the OC/ON niche, for the indicated ligand–receptor pairs. Each dot represents one case from the NR, RD, or RI groups. Spearman rank correlation coefficients (r) and p-values are shown in each plot.

### Supplementary Figure 8

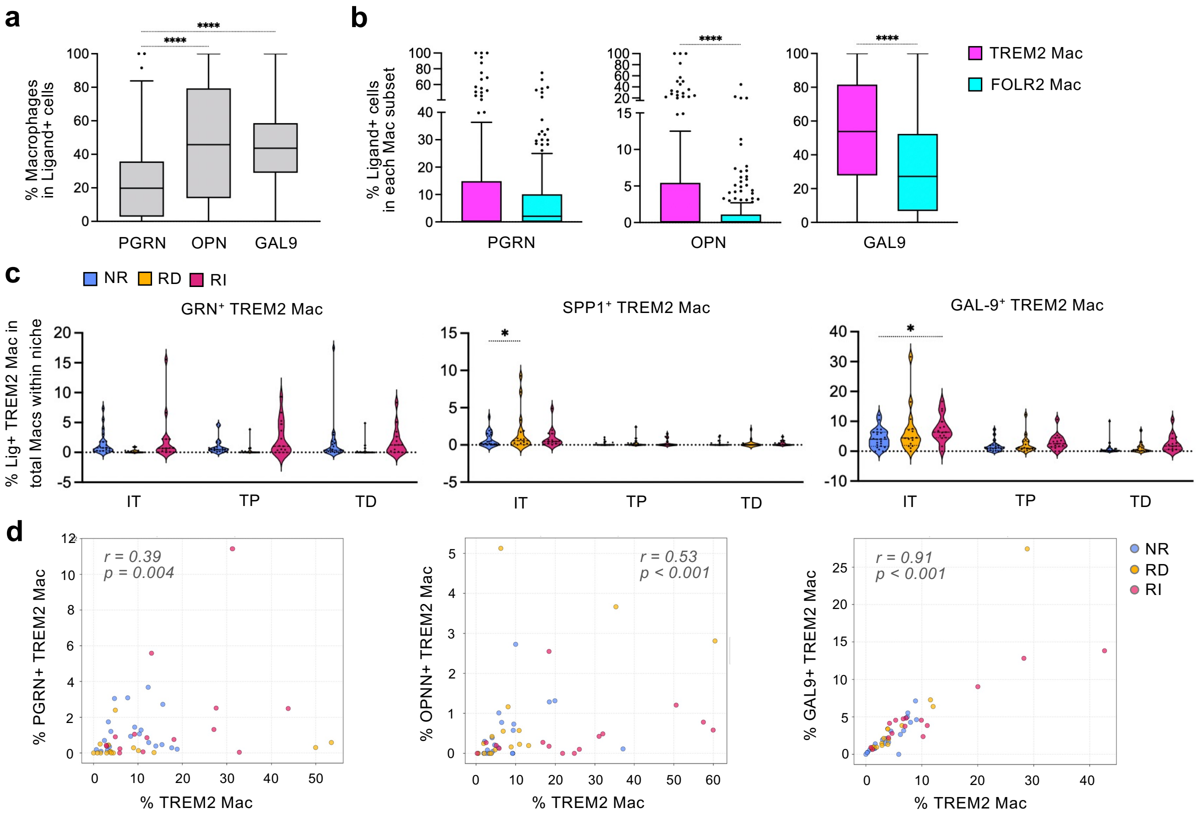


**Supplementary Fig. 8 Spatial mapping of candidate ligand proteins in DCIS. a.** Tukey box plots showing the proportions of macrophages among total ligand-positive cells across cores and niches. Statistical differences across ligands were assessed using Kruskal–Wallis tests followed by Dunn's multiple-comparison test (****$p < 0.0001$). **b.** Tukey box plots showing the proportions of ligand-positive cells among total TREM2+ or FOLR2+ macrophages across cores and niches. Statistical differences between macrophage subsets were assessed using Mann–Whitney test (****$p < 0.0001$). **c.** Violin plots showing the proportion of indicated ligand-positive populations among all TREM2+ macrophages within each niche across patient groups (NR, RD, and RI). Each dot represents one case. Statistical differences among patient groups were assessed by two-way ANOVA followed by Tukey's multiple-comparisons test (*$p < 0.05$). **d.** Correlations between the proportion of ligand-positive TREM2$^{+}$ Macs (y-axis) and the proportion of TREM2+ Macs (x-axis), both calculated among all macrophages across niches. Each dot represents one case from the NR, RD, or RI groups. Spearman rank correlation coefficients (r) and p-values are shown in each plot. Mac, macrophage; PGRN, progranulin; OPN, osteopontin; GAL9, galectin 9; IT, intra-tumoral niche; TP, tumor-proximal niche; TD, tumor-distal niche; NR, no recurrence; RD, recurrent DCIS; RI, recurrent IBC.

#### Supplementary Table 1

| Sample ID | Grade | Age | ER status | PR status | Margin width |
|---|---|---|---|---|---|
| DCIS-1 | Low | 47 | + | N/A | ≧ 1 mm |
| DCIS-2 | Low | 47 | + | N/A | ≧ 1 mm |
| DCIS-3 | Intermediate | 68 | + | N/A | ≧ 1 mm |
| DCIS-4 | Intermediate | 59 | + | N/A | < 1 mm |
| DCIS-5 | High | 53 | – | N/A | < 1 mm |
| DCIS-6 | High | 42 | – | N/A | < 1 mm |
| DCIS-7 | High | 64 | + | N/A | < 1 mm |
| DCIS-8 | High | 64 | + | N/A | < 1 mm |
| IBC-1 | Invasive | 63 | + | + | ≧ 1 mm |
| IBC-2 | Invasive | 49 | + | + | Positive (0 mm) |
| IBC-3 | Invasive | 64 | + | + | < 1 mm |
| IBC-4 | Invasive | 53 | + | + | ≧ 1 mm |

**Supplementary Table 1 Description of samples used for snRNA-seq.** Retrospective cohort of patients diagnosed with lo/intermediate-grade DCIS (N = 4), high-grade DCIS (N = 4) or IBC (N = 4) after surgery (Cohort-1). snRNA-seq, single-nuclei RNA sequencing; ER, estrogen receptor; PR, progesterone receptor; N/A, not available.

#### Supplementary Table 2

| | No recurrence (N=21) | Recurrent DCIS (N=14) | Recurrent IBC (N=14) |
|---|---|---|---|
| **Age at diagnosis** | | | |
| Median | 59 | 57 | 55 |
| Mean (± SD) | 59 (± 9.8) | 58 (± 6.0) | 57 (± 10.2) |
| **Grade** | | | |
| 1 | 0 [0%] | 1 [7.1%] | 0 [0%] |
| 2 | 5 [23.8%] | 4 [28.6%] | 4 [28.6%] |
| 3 | 16 [76.2%] | 9 [64.3%] | 10 [71.4%] |
| **Tumor size (mm)** | | | |
| Median | 14 | 26.5 | 20 |
| Mean (± SD) | 18 (± 11.3) | 27 (± 8.5) | 22 (± 9.0) |
| **Marker status** | | | |
| ER (+) | 18 [85.7%] | 11 [78.6%] | 9 [64.3%] |
| ER (–) | 0 [0%] | 1 [7.1%] | 1 [7.1%] |
| N/A | 3 [14.3%] | 2 [14.3%] | 4 [28.6%] |
| **Surgery** | | | |
| BCS | 20 [95.2%] | 13 [92.9%] | 14 [100%] |
| BCS + ANS | 1 [4.8%] | 1 [7.1%] | 0 [0%] |
| **Margin width** | | | |
| ≦ 1 mm | 5 [23.8%] | 2 [14.3%] | 2 [14.3%] |
| > 1 mm | 16 [76.2%] | 12 [85.7%] | 12 [85.7%] |
| **Endocrine therapy** | | | |
| None | 20 [95.2%] | 13 [92.9%] | 13 [92.9%] |
| Adjuvant | 1 [4.8%] | 0 [0%] | 1 [7.1%] |
| Neo-adjuvant | 0 [0%] | 1 [7.1%] | 0 [0%] |
| **Radiotherapy** | | | |
| None | 5 [23.8%] | 3 [21.4%] | 2 [14.3%] |
| Boost only | 0 [0%] | 2 [14.3%] | 2 [14.3%] |
| Standard | 16 [76.2%] | 9 [64.3%] | 9 [71.4%] |
| **Time to recurrence (years)*** | | | |
| Median | 8.1 | 2.6 | 2.5 |
| Mean (± SD) | 7.9 (± 3.0) | 4.0 (± 2.8) | 3.7 (± 2.2) |

*To end of follow-up for no recurrence

**Supplementary Table 2 Description of cohorts used for spatial transcriptomics.** Retrospective cohort of DCIS patients who did not recur tumors (No recurrence, N = 21), recurred DCIS without IBC (Recurrent DCIS, N = 14) or recurred IBC (Recurrent IBC, N = 14) during follow-up (Cohort-2). ER, estrogen receptor; N/A, not available; BCS, breast-conserving surgery: ANS, axillary node surgery.

#### Supplementary Table 3

| No recurrence (NR) | | | Recurrenct DCIS (RD) | | | Recurrent IBC (RI) | | |
|---|---|---|---|---|---|---|---|---|
| Patient ID | # Analyzable Cores | # Analyzable ROIs | Patient ID | # Analyzable Cores | # Analyzable ROIs | Patient ID | # Analyzable Cores | # Analyzable ROIs |
| SR2119-038 | 4 | 14 | SR2119-017 | 0 | 0 | SR2119-001 | 1 | 4 |
| SR2119-039 | 5 | 19 | SR2119-018 | 1 | 2 | SR2119-002 | 5 | 28 |
| SR2119-040 | 4 | 14 | SR2119-019 | 5 | 22 | SR2119-003 | 1 | 2 |
| SR2119-041 | 3 | 11 | SR2119-020 | 7 | 24 | SR2119-004 | 7 | 31 |
| SR2119-042 | 5 | 29 | SR2119-021 | 6 | 23 | SR2119-005 | 2 | 4 |
| SR2119-043 | 3 | 11 | SR2119-022 | 2 | 5 | SR2119-006 | 4 | 19 |
| SR2119-044 | 4 | 7 | SR2119-023 | 2 | 3 | SR2119-007 | 2 | 7 |
| SR2119-045 | 5 | 27 | SR2119-024 | 3 | 9 | SR2119-008 | 1 | 2 |
| SR2119-046 | 4 | 15 | SR2119-025 | 1 | 2 | SR2119-009 | 6 | 28 |
| SR2119-047 | 3 | 7 | SR2119-026 | 5 | 15 | SR2119-010 | 4 | 13 |
| SR2119-048 | 5 | 15 | SR2119-027 | 5 | 21 | SR2119-011 | 4 | 11 |
| SR2119-049 | 5 | 27 | SR2119-028 | 1 | 2 | SR2119-012 | 3 | 10 |
| SR2119-050 | 1 | 1 | SR2119-029 | 3 | 13 | SR2119-013 | 0 | 0 |
| SR2119-051 | 3 | 15 | SR2119-030 | 1 | 2 | SR2119-014 | 5 | 20 |
| SR2119-052 | 3 | 21 | SR2119-031 | 0 | 0 | SR2119-016 | 3 | 11 |
| SR2119-053 | 0 | 0 | SR2119-032 | 3 | 8 | | | |
| SR2119-054 | 6 | 30 | | | | | | |
| SR2119-055 | 2 | 8 | | | | | | |
| SR2119-056 | 5 | 23 | | | | | | |
| SR2119-057 | 2 | 9 | | | | | | |
| SR2119-058 | 5 | 19 | | | | | | |
| SR2119-059 | 5 | 26 | | | | | | |
| Total | 82 | 348 | Total | 45 | 151 | Total | 48 | 190 |
| Median | 4 | 15 | Median | 2.5 | 6.5 | Median | 3 | 11 |
| Mean | 3.7 | 15.8 | Mean | 2.8 | 9.4 | Mean | 3.2 | 12.7 |

**Supplementary Table 3 Number of cores and regions of interests used for spatial transcriptomics.** Each case included 1-6 cores, comprising 1–8 ROIs. ROIs were excluded if they met any of the following criteria: fewer than 200 transcripts per cell, partial or complete tissue detachment, absence of tumor nests (epithelial cell clusters), or insufficient cellularity (no or very few cells). Cores without any eligible ROIs were excluded from analysis. ROIs, regions of interest.

### Supplementary Table 4

**a** **Up in TREM2 Mac**: 524genes

| Gene | log2FC | padj |
|---|---|---|
| CHRNA1 | 7.01 | 1.7E-07 |
| SPOCD1 | 6.40 | 9.4E-08 |
| CHIT1 | 5.84 | 4.3E-10 |
| SHISAL2A | 5.30 | 7.1E-11 |
| TM4SF19 | 5.24 | 5.8E-11 |
| CHI3L1 | 5.02 | 2.7E-11 |
| HAMP | 4.97 | 2.3E-10 |
| HES2 | 4.83 | 3.1E-05 |
| CD207 | 4.76 | 1.1E-03 |
| ALKAL2 | 4.74 | 2.7E-05 |
| MMP7 | 4.70 | 4.7E-06 |
| NMRK2 | 4.70 | 6.7E-04 |
| DCSTAMP | 4.67 | 5.2E-06 |
| RTN4R | 4.58 | 1.3E-08 |
| LILRA4 | 4.39 | 4.5E-08 |
| NRIP3 | 4.39 | 3.1E-09 |
| TMEM52B | 4.37 | 6.6E-08 |
| TGM2 | 4.32 | 9.8E-18 |
| SYNDIG1 | 4.23 | 2.7E-06 |
| GP2 | 4.13 | 2.8E-02 |
| KCP | 4.13 | 1.2E-10 |
| CNIH3 | 4.12 | 1.6E-08 |
| NIPAL4 | 4.10 | 8.5E-03 |
| CLEC17A | 4.05 | 7.3E-06 |
| MS4A6E | 4.03 | 2.3E-03 |
| APOC2 | 4.02 | 1.4E-08 |
| SPSB1 | 4.02 | 1.3E-20 |
| ITGB3 | 4.00 | 9.2E-17 |
| TMEM255A | 3.98 | 2.3E-09 |
| SLAMF7 | 3.96 | 2.6E-18 |
| CCL22 | 3.94 | 1.1E-04 |
| FABP4 | 3.91 | 1.4E-03 |
| APOC1 | 3.86 | 5.1E-13 |
| ADGRG5 | 3.84 | 9.8E-12 |
| KCNQ3 | 3.82 | 3.0E-10 |
| TNFSF14 | 3.82 | 2.4E-09 |
| CES1 | 3.81 | 1.2E-04 |
| ITGA3 | 3.81 | 2.0E-16 |
| PADI2 | 3.81 | 1.1E-14 |
| MCOLN3 | 3.77 | 4.1E-12 |
| PKD2L1 | 3.75 | 1.0E-06 |
| CD52 | 3.74 | 6.8E-13 |
| GCHFR | 3.72 | 7.6E-13 |
| HTRA4 | 3.69 | 3.6E-14 |
| NCAPH | 3.69 | 5.4E-04 |
| SALL1 | 3.67 | 5.7E-04 |
| ADGRG1 | 3.67 | 1.2E-08 |
| CLDN1 | 3.66 | 2.3E-10 |
| HSD11B1 | 3.63 | 9.0E-03 |
| MMP8 | 3.61 | 5.9E-03 |
| SLC12A3 | 3.61 | 6.7E-06 |
| ATP6V0D2 | 3.58 | 8.6E-09 |
| GDF15 | 3.57 | 1.9E-10 |
| CSF1 | 3.57 | 7.6E-13 |
| SULT1C2 | 3.56 | 6.7E-05 |
| HRK | 3.56 | 3.9E-03 |
| PGBD5 | 3.54 | 1.3E-14 |
| PNPLA3 | 3.53 | 6.6E-08 |
| OLR1 | 3.50 | 1.1E-10 |
| MELTF | 3.50 | 7.4E-10 |
| SIGLEC8 | 3.42 | 4.6E-10 |
| ADGRE3 | 3.41 | 5.5E-04 |
| FCGBP | 3.41 | 4.5E-10 |
| MMP12 | 3.39 | 5.1E-04 |
| MATK | 3.39 | 5.2E-07 |
| DNASE2B | 3.38 | 8.0E-07 |
| AQP9 | 3.36 | 1.8E-07 |
| TMEM156 | 3.36 | 3.5E-07 |
| CAMK1G | 3.36 | 2.1E-02 |
| OLFML3 | 3.35 | 5.9E-15 |
| NCS1 | 3.30 | 4.2E-07 |
| LPL | 3.30 | 5.6E-09 |
| IL1A | 3.30 | 5.2E-03 |
| TLR10 | 3.27 | 1.2E-12 |
| SPN | 3.27 | 5.6E-36 |
| CLEC4F | 3.25 | 7.2E-03 |
| IFNLR1 | 3.23 | 2.0E-10 |
| MREG | 3.22 | 5.4E-13 |
| CASQ1 | 3.22 | 1.6E-03 |
| SCIN | 3.21 | 2.3E-07 |
| ACOT4 | 3.21 | 5.4E-09 |
| EML1 | 3.21 | 6.8E-06 |
| AFAP1L1 | 3.20 | 1.5E-05 |
| C3 | 3.20 | 1.2E-09 |
| CD6 | 3.19 | 1.3E-06 |
| MACC1 | 3.18 | 5.4E-15 |
| MYLK2 | 3.17 | 3.6E-02 |
| CPEB1 | 3.17 | 3.0E-03 |

| Gene | log2FC | padj |
|---|---|---|
| ASPHD1 | 3.16 | 3.7E-03 |
| RASL11B | 3.15 | 3.6E-02 |
| IL1B | 3.14 | 1.6E-05 |
| GPC4 | 3.14 | 3.3E-06 |
| WNT5A | 3.14 | 2.4E-06 |
| SLC2A1 | 3.13 | 1.7E-09 |
| TNFRSF9 | 3.12 | 4.2E-03 |
| TDRD9 | 3.12 | 1.0E-04 |
| RARRES1 | 3.11 | 3.3E-11 |
| ABCB5 | 3.10 | 2.9E-04 |
| BEAN1 | 3.10 | 3.2E-03 |
| CAPG | 3.09 | 1.2E-17 |
| CP | 3.09 | 6.0E-04 |
| SDS | 3.09 | 1.4E-08 |
| HPGD | 3.08 | 2.3E-03 |
| PLEK2 | 3.08 | 1.8E-06 |
| GPR143 | 3.07 | 9.2E-03 |
| CCL20 | 3.06 | 1.6E-03 |
| CADM1 | 3.06 | 1.1E-14 |
| CHRNA6 | 3.05 | 4.4E-04 |
| LPAR3 | 3.05 | 2.8E-02 |
| JAKMIP2 | 3.05 | 7.2E-03 |
| ELAVL4 | 3.04 | 2.3E-05 |
| ADD2 | 3.04 | 3.8E-05 |
| PLAC8 | 3.03 | 5.5E-08 |
| MANSC4 | 3.02 | 3.7E-02 |
| USP2 | 3.01 | 1.5E-04 |
| GOLGA7B | 3.00 | 5.6E-04 |
| FAM149A | 2.98 | 5.8E-07 |
| FCMR | 2.97 | 2.2E-10 |
| GALNT12 | 2.97 | 8.7E-10 |
| ILDR2 | 2.97 | 2.0E-02 |
| MYOZ1 | 2.97 | 3.8E-03 |
| ADAMTSL2 | 2.97 | 1.2E-03 |
| IFI27 | 2.96 | 6.7E-04 |
| TENM4 | 2.96 | 1.6E-03 |
| HK2 | 2.95 | 5.6E-19 |
| LY9 | 2.94 | 2.2E-08 |
| CLEC9A | 2.94 | 3.2E-04 |
| RASGRF1 | 2.94 | 4.3E-05 |
| SPNS3 | 2.93 | 2.0E-05 |
| PROCR | 2.93 | 2.1E-08 |
| BCL2A1 | 2.92 | 2.1E-15 |
| PROC | 2.92 | 2.4E-03 |
| GALNTL6 | 2.91 | 9.3E-02 |
| CX3CR1 | 2.89 | 1.9E-06 |
| RHOF | 2.88 | 6.1E-09 |
| DOCK3 | 2.87 | 6.7E-06 |
| CD9 | 2.87 | 2.0E-29 |
| RBP4 | 2.85 | 1.4E-03 |
| TSPAN12 | 2.84 | 9.3E-04 |
| CA2 | 2.84 | 2.0E-10 |
| MME | 2.83 | 4.3E-06 |
| CCR6 | 2.83 | 8.9E-05 |
| SPNS2 | 2.83 | 3.4E-07 |
| SYT6 | 2.83 | 1.7E-06 |
| BHLHE41 | 2.82 | 5.2E-31 |
| ADAM8 | 2.81 | 2.0E-12 |
| TREM2 | 2.81 | 4.2E-19 |
| FAIM | 2.80 | 4.6E-11 |
| MCF2L | 2.80 | 2.1E-06 |
| SLC6A7 | 2.79 | 5.3E-02 |
| ADAMTS15 | 2.78 | 1.3E-03 |
| UNC5B | 2.78 | 1.3E-06 |
| SLAMF9 | 2.78 | 1.6E-03 |
| MRC2 | 2.77 | 2.9E-14 |
| PHLDA3 | 2.76 | 2.3E-07 |
| KLK4 | 2.76 | 1.1E-02 |
| CAMP | 2.76 | 3.0E-02 |
| CD300LB | 2.76 | 9.1E-07 |
| APOBEC3A | 2.75 | 5.9E-02 |
| SMIM24 | 2.75 | 7.1E-02 |
| LINC02898 | 2.75 | 4.1E-02 |
| ADGRB3 | 2.71 | 1.7E-02 |
| ALDH1A2 | 2.70 | 2.4E-03 |
| PARM1 | 2.69 | 7.7E-06 |
| YPEL4 | 2.67 | 4.2E-05 |
| RGCC | 2.67 | 5.2E-12 |
| NDRG4 | 2.64 | 2.0E-02 |
| MCEMP1 | 2.64 | 4.4E-02 |
| SLC28A3 | 2.62 | 1.2E-03 |
| GSDME | 2.61 | 3.6E-13 |
| AGMAT | 2.61 | 2.0E-02 |
| DNAJC5B | 2.61 | 3.5E-06 |
| PAQR5 | 2.60 | 5.6E-05 |
| PLD4 | 2.58 | 6.5E-06 |
| FLT1 | 2.58 | 5.6E-04 |
| CYP27A1 | 2.57 | 2.3E-10 |

| Gene | log2FC | padj |
|---|---|---|
| C15orf48 | 2.56 | 8.8E-08 |
| SLC7A5 | 2.55 | 3.4E-14 |
| CD22 | 2.54 | 2.3E-08 |
| SLC47A1 | 2.54 | 8.6E-06 |
| TMEM266 | 2.53 | 2.4E-05 |
| G0S2 | 2.53 | 7.7E-04 |
| SIGLEC10 | 2.52 | 1.7E-07 |
| FGR | 2.51 | 1.1E-13 |
| STUM | 2.51 | 7.9E-03 |
| CDH23 | 2.50 | 2.1E-10 |
| ALOX15B | 2.50 | 4.6E-05 |
| PRAM1 | 2.50 | 1.8E-22 |
| NECTIN4 | 2.50 | 6.2E-06 |
| CXCR2 | 2.49 | 1.2E-02 |
| DYNLT2B | 2.49 | 6.3E-05 |
| APOC4 | 2.47 | 8.0E-04 |
| MSX2 | 2.47 | 5.5E-02 |
| ANOS1 | 2.47 | 2.3E-07 |
| BAALC | 2.46 | 7.3E-02 |
| RHOH | 2.46 | 2.3E-09 |
| LHX2 | 2.46 | 9.5E-02 |
| ICAM3 | 2.46 | 2.1E-06 |
| ENO2 | 2.46 | 6.3E-07 |
| TREML1 | 2.46 | 1.3E-08 |
| MFSD2A | 2.45 | 3.0E-08 |
| MOSPD1 | 2.45 | 1.3E-10 |
| MEP1A | 2.45 | 3.1E-02 |
| NUPR1 | 2.45 | 4.9E-09 |
| LSP1 | 2.44 | 1.2E-19 |
| FBP1 | 2.44 | 3.2E-09 |
| ALCAM | 2.44 | 1.3E-16 |
| PLVAP | 2.43 | 3.3E-09 |
| SCD | 2.43 | 1.1E-10 |
| EDA2R | 2.42 | 1.4E-03 |
| CCDC148 | 2.42 | 5.7E-02 |
| RBP1 | 2.42 | 1.8E-04 |
| AZIN2 | 2.41 | 3.9E-06 |
| KIF26B | 2.41 | 1.8E-05 |
| CCL7 | 2.41 | 4.1E-02 |
| CD226 | 2.41 | 1.9E-06 |
| CD109 | 2.41 | 2.8E-12 |
| NPFFR1 | 2.40 | 9.9E-02 |
| ADTRP | 2.40 | 4.7E-04 |
| SLC1A4 | 2.40 | 2.3E-14 |
| GLDN | 2.39 | 1.8E-02 |
| FABP5 | 2.39 | 7.0E-05 |
| CHST2 | 2.39 | 1.2E-04 |
| MOGAT1 | 2.39 | 9.0E-02 |
| ADCY3 | 2.38 | 3.1E-18 |
| DHDH | 2.38 | 1.0E-03 |
| RAI14 | 2.36 | 2.2E-03 |
| KEL | 2.35 | 6.4E-04 |
| GNAO1 | 2.35 | 8.8E-03 |
| TNFAIP8L3 | 2.35 | 4.3E-03 |
| TSPAN10 | 2.33 | 2.1E-02 |
| APOE | 2.33 | 4.3E-11 |
| PKIB | 2.33 | 1.4E-04 |
| RGS13 | 2.33 | 1.9E-02 |
| MLXIPL | 2.33 | 2.1E-04 |
| CLEC5A | 2.32 | 2.1E-04 |
| GPR18 | 2.32 | 6.0E-03 |
| RAB36 | 2.32 | 1.8E-02 |
| DDIT4L | 2.31 | 9.9E-02 |
| VASH1 | 2.30 | 1.0E-03 |
| SLC2A5 | 2.29 | 4.0E-05 |
| APOBR | 2.29 | 2.9E-15 |
| MYBPH | 2.28 | 9.4E-02 |
| TMEM217 | 2.28 | 2.0E-02 |
| CD1E | 2.27 | 1.8E-02 |
| KCNA3 | 2.27 | 3.0E-04 |
| TRO | 2.26 | 8.2E-04 |
| CPA6 | 2.26 | 1.1E-01 |
| ABI3 | 2.26 | 6.9E-12 |
| SIRPB1 | 2.24 | 3.1E-07 |
| TSKU | 2.24 | 5.0E-04 |
| NUAK1 | 2.23 | 1.7E-06 |
| CAMK2B | 2.23 | 3.3E-02 |
| TMEM44 | 2.23 | 2.4E-06 |
| CLMP | 2.23 | 3.0E-04 |
| DACT3 | 2.20 | 1.6E-04 |
| SDK2 | 2.20 | 3.6E-02 |
| ANGPTL6 | 2.19 | 1.8E-03 |
| FDX1 | 2.19 | 2.8E-12 |
| MLPH | 2.18 | 5.5E-07 |
| OLFM2 | 2.18 | 4.1E-03 |
| CD48 | 2.17 | 3.6E-10 |
| PBX4 | 2.17 | 5.8E-03 |
| PTPN7 | 2.16 | 1.4E-09 |

| Gene | log2FC | padj |
|---|---|---|
| COL8A2 | 2.16 | 9.2E-17 |
| ACE | 2.16 | 2.7E-04 |
| ITGB7 | 2.16 | 3.7E-07 |
| PRR15 | 2.14 | 6.5E-03 |
| CD82 | 2.14 | 4.8E-17 |
| PDGFA | 2.14 | 4.0E-06 |
| CST7 | 2.14 | 4.5E-03 |
| NUP210 | 2.13 | 4.2E-15 |
| IPCEF1 | 2.13 | 1.9E-04 |
| NKD1 | 2.12 | 2.9E-02 |
| DENND2B | 2.11 | 2.5E-15 |
| IL17B | 2.10 | 7.5E-02 |
| SORL1 | 2.10 | 1.6E-09 |
| CYGB | 2.10 | 4.3E-04 |
| RFLNB | 2.09 | 1.2E-03 |
| QPCT | 2.09 | 6.1E-06 |
| CSTB | 2.08 | 2.1E-08 |
| CXADR | 2.08 | 7.5E-03 |
| RUNX3 | 2.08 | 9.2E-08 |
| OR52K1 | 2.07 | 2.4E-02 |
| FAIM2 | 2.07 | 7.3E-02 |
| RAB11FIP4 | 2.07 | 2.5E-06 |
| SARDH | 2.07 | 1.6E-02 |
| DCLK2 | 2.07 | 1.7E-03 |
| SH3D21 | 2.07 | 4.6E-07 |
| SEZ6L2 | 2.07 | 1.8E-03 |
| SDC4 | 2.07 | 2.3E-12 |
| CD3D | 2.06 | 3.3E-02 |
| ENPP1 | 2.06 | 2.2E-04 |
| WARS1 | 2.06 | 5.8E-11 |
| SPATA13 | 2.06 | 5.3E-13 |
| GNGT2 | 2.06 | 4.5E-07 |
| MLLT11 | 2.05 | 1.6E-02 |
| RAB6B | 2.05 | 3.8E-03 |
| RGS1 | 2.04 | 8.2E-04 |
| AGRP | 2.04 | 5.3E-02 |
| MTHFS | 2.04 | 1.2E-10 |
| CARD11 | 2.03 | 7.6E-06 |
| PRSS22 | 2.03 | 1.1E-01 |
| FCRLB | 2.03 | 2.1E-02 |
| SLC6A12 | 2.02 | 7.0E-05 |
| DPP4 | 2.02 | 1.5E-03 |
| PLA2G7 | 2.02 | 9.2E-09 |
| CLIC3 | 2.02 | 2.7E-02 |
| IL1RN | 2.02 | 1.6E-02 |
| DNAJB5 | 2.01 | 1.4E-04 |
| TLN2 | 2.01 | 6.9E-07 |
| ACTN1 | 2.00 | 1.6E-19 |
| IFITM10 | 2.00 | 8.1E-16 |
| CENPV | 2.00 | 9.2E-03 |
| FAM167A | 1.99 | 5.2E-02 |
| RASGEF1A | 1.98 | 3.5E-02 |
| RCAN3 | 1.98 | 2.5E-14 |
| CKB | 1.98 | 2.9E-03 |
| YWHAH | 1.97 | 5.2E-12 |
| ADCY1 | 1.97 | 6.9E-02 |
| SNTB1 | 1.97 | 4.5E-06 |
| SORT1 | 1.97 | 5.2E-24 |
| TIGD4 | 1.97 | 1.3E-02 |
| CA11 | 1.96 | 5.2E-05 |
| SLC2A6 | 1.96 | 2.7E-03 |
| PI15 | 1.95 | 9.4E-02 |
| GAL3ST4 | 1.95 | 1.2E-06 |
| LGALS3 | 1.94 | 2.4E-11 |
| IFI6 | 1.94 | 9.4E-04 |
| LGALSL | 1.93 | 3.7E-05 |
| SEMA3C | 1.93 | 9.1E-08 |
| ITGB8 | 1.93 | 6.7E-04 |
| DYRK3 | 1.92 | 3.3E-02 |
| DCDC2 | 1.91 | 1.6E-02 |
| NEURL3 | 1.91 | 1.1E-02 |
| P2RX5 | 1.91 | 2.3E-02 |
| SDC2 | 1.91 | 4.3E-04 |
| GPAT3 | 1.91 | 5.7E-08 |
| PC | 1.91 | 1.1E-06 |
| FGF20 | 1.90 | 1.7E-02 |
| KCNK13 | 1.90 | 7.3E-08 |
| DHCR24 | 1.90 | 1.2E-04 |
| FJX1 | 1.90 | 5.9E-02 |
| TOP2A | 1.90 | 4.7E-02 |
| CTSV | 1.89 | 4.2E-03 |
| CDH1 | 1.89 | 2.5E-04 |
| CDKN2A | 1.89 | 1.3E-03 |
| ALPK3 | 1.89 | 4.8E-05 |
| NR1H3 | 1.88 | 4.9E-10 |
| LIMK1 | 1.87 | 6.6E-21 |
| OR52K2 | 1.87 | 9.7E-03 |
| PALM2AKAP2 | 1.87 | 3.6E-14 |

| Gene | log2FC | padj |
|---|---|---|
| LIPA | 1.86 | 3.7E-07 |
| ACP5 | 1.86 | 7.6E-08 |
| GPR157 | 1.86 | 1.1E-04 |
| ISG15 | 1.86 | 1.3E-02 |
| RGS16 | 1.85 | 4.6E-03 |
| SPATA12 | 1.85 | 4.2E-02 |
| WIPF3 | 1.85 | 8.4E-02 |
| OPTN | 1.85 | 6.2E-04 |
| RNF128 | 1.85 | 1.0E-01 |
| ID2 | 1.85 | 1.2E-23 |
| PPA1 | 1.84 | 5.0E-08 |
| TBC1D1 | 1.84 | 1.6E-08 |
| ATP6V1F | 1.84 | 5.1E-12 |
| STRBP | 1.83 | 3.8E-11 |
| B4GALT5 | 1.83 | 1.2E-06 |
| GMPR | 1.83 | 9.4E-03 |
| ZBP1 | 1.83 | 1.0E-02 |
| PPP1R1B | 1.83 | 9.8E-02 |
| PPP1R16B | 1.83 | 1.1E-02 |
| SERINC2 | 1.83 | 1.4E-05 |
| GRIN3A | 1.83 | 8.3E-02 |
| SLC26A11 | 1.83 | 2.0E-06 |
| SDSL | 1.82 | 1.3E-08 |
| TRIB3 | 1.82 | 3.2E-02 |
| ZNF385A | 1.82 | 8.9E-21 |
| ACSM3 | 1.82 | 4.1E-02 |
| MARCHF3 | 1.82 | 3.4E-04 |
| AMZ1 | 1.81 | 6.5E-02 |
| SLC6A16 | 1.81 | 7.7E-03 |
| SHC3 | 1.80 | 6.4E-02 |
| TMEM163 | 1.80 | 2.2E-02 |
| DNASE2 | 1.80 | 7.2E-09 |
| GPNMB | 1.80 | 1.3E-10 |
| KIF23 | 1.80 | 6.0E-02 |
| ADGRE2 | 1.80 | 2.4E-06 |
| IFITM1 | 1.79 | 5.8E-02 |
| SPP1 | 1.79 | 9.7E-03 |
| MAFF | 1.79 | 1.5E-04 |
| FGD5 | 1.78 | 6.5E-08 |
| RALA | 1.78 | 1.0E-10 |
| BMERB1 | 1.78 | 4.8E-04 |
| ST14 | 1.78 | 2.6E-11 |
| SLC35F2 | 1.78 | 3.0E-03 |
| FABP3 | 1.77 | 5.0E-07 |
| MAP9 | 1.77 | 1.3E-02 |
| CCR7 | 1.76 | 4.2E-02 |
| MYO1D | 1.76 | 3.4E-07 |
| MICAL1 | 1.76 | 8.1E-25 |
| CD300LF | 1.76 | 5.0E-10 |
| PLA2G2D | 1.75 | 5.1E-02 |
| FOXM1 | 1.75 | 8.6E-02 |
| HSD3B7 | 1.75 | 2.3E-07 |
| PLXNC1 | 1.75 | 3.0E-07 |
| SOX13 | 1.75 | 3.7E-02 |
| SPRING1 | 1.75 | 1.0E-04 |
| PPIAL4G | 1.75 | 9.2E-02 |
| TMCC3 | 1.75 | 6.6E-06 |
| OR3A1 | 1.74 | 9.8E-02 |
| BOLA3 | 1.74 | 2.7E-03 |
| MRAS | 1.74 | 5.6E-07 |
| PRSS21 | 1.74 | 8.3E-03 |
| TLR3 | 1.73 | 1.3E-05 |
| TMEM154 | 1.73 | 9.4E-04 |
| CCR2 | 1.73 | 9.4E-03 |
| CYB561A3 | 1.73 | 3.5E-09 |
| C12orf75 | 1.72 | 6.3E-03 |
| BIRC7 | 1.72 | 4.9E-02 |
| PDPN | 1.72 | 8.1E-03 |
| SEMA7A | 1.71 | 2.2E-07 |
| SOAT1 | 1.71 | 4.3E-13 |
| S100A11 | 1.71 | 1.0E-07 |
| BLOC1S2 | 1.71 | 2.2E-07 |
| ST8SIA6 | 1.70 | 6.5E-02 |
| CELSR1 | 1.70 | 1.1E-04 |
| ABCG2 | 1.70 | 2.2E-03 |
| SLC9A7 | 1.70 | 5.4E-08 |
| MCOLN2 | 1.70 | 1.5E-04 |
| GNAI1 | 1.70 | 3.1E-04 |
| PAPLN | 1.69 | 1.9E-05 |
| ADAMDEC1 | 1.69 | 3.3E-05 |
| JAM2 | 1.69 | 3.4E-03 |
| MAP2 | 1.69 | 8.5E-02 |
| GM2A | 1.67 | 1.7E-10 |
| C3orf52 | 1.67 | 1.0E-01 |
| EPB41L2 | 1.67 | 1.5E-19 |
| GAREM1 | 1.66 | 1.3E-02 |
| TIMP3 | 1.65 | 7.4E-03 |
| TNFRSF12A | 1.65 | 6.0E-07 |

| Gene | log2FC | padj |
|---|---|---|
| PRKCH | 1.65 | 1.0E-05 |
| MAMLD1 | 1.65 | 1.6E-02 |
| TMEM231 | 1.65 | 1.2E-02 |
| COL27A1 | 1.64 | 3.8E-03 |
| RASSF6 | 1.64 | 3.4E-02 |
| FAM111B | 1.64 | 6.9E-02 |
| RAB42 | 1.64 | 7.2E-06 |
| PLP2 | 1.64 | 2.3E-12 |
| KALRN | 1.63 | 4.3E-02 |
| MYO10 | 1.63 | 4.6E-05 |
| TBC1D10C | 1.63 | 5.9E-09 |
| BBLN | 1.63 | 6.0E-07 |
| ATP9A | 1.62 | 7.8E-04 |
| AXDND1 | 1.62 | 1.2E-01 |
| TPRG1 | 1.61 | 1.0E-03 |
| RAB7B | 1.61 | 1.9E-04 |
| SHOX2 | 1.61 | 3.6E-02 |
| HSPB8 | 1.61 | 2.5E-02 |
| CD80 | 1.61 | 2.4E-04 |
| CD81 | 1.61 | 8.2E-12 |
| BIRC5 | 1.61 | 9.7E-02 |
| SLC29A1 | 1.61 | 5.3E-11 |
| FBXL2 | 1.61 | 8.3E-02 |
| GPX7 | 1.61 | 3.7E-03 |
| HCAR1 | 1.60 | 7.2E-02 |
| NDRG2 | 1.60 | 2.5E-04 |
| RASSF8 | 1.60 | 9.6E-06 |
| SMAD7 | 1.60 | 2.6E-06 |
| SEMA3A | 1.60 | 2.6E-02 |
| PLEKHF1 | 1.60 | 6.4E-03 |
| CD101 | 1.60 | 1.6E-02 |
| PALD1 | 1.59 | 7.1E-04 |
| EDARADD | 1.59 | 1.1E-01 |
| DNAI3 | 1.59 | 9.9E-02 |
| CD1C | 1.59 | 4.0E-02 |
| UPP1 | 1.59 | 3.3E-09 |
| LMO7 | 1.59 | 7.6E-04 |
| SH3BP4 | 1.59 | 3.5E-04 |
| GAPT | 1.59 | 7.2E-06 |
| ANXA2 | 1.58 | 2.1E-05 |
| SERPINF1 | 1.58 | 9.6E-07 |
| CCRL2 | 1.58 | 2.2E-06 |
| CTSD | 1.57 | 3.9E-07 |
| GPR84 | 1.57 | 2.5E-04 |
| AHNAK2 | 1.57 | 1.2E-02 |
| OXCT1 | 1.57 | 7.2E-04 |
| CTH | 1.56 | 4.6E-02 |
| PGAM1 | 1.56 | 1.1E-03 |
| NCEH1 | 1.56 | 9.7E-08 |
| STRIP2 | 1.56 | 2.0E-02 |
| PLAUR | 1.56 | 2.6E-13 |
| JPT1 | 1.56 | 4.4E-06 |
| RSAD2 | 1.56 | 6.3E-02 |
| SPINT1 | 1.55 | 4.9E-10 |
| CORO2A | 1.55 | 3.8E-05 |
| SOCS6 | 1.55 | 1.6E-13 |
| SEL1L3 | 1.55 | 2.3E-11 |
| P2RY12 | 1.55 | 6.3E-04 |
| SP140 | 1.55 | 2.8E-07 |
| SMAD6 | 1.55 | 2.1E-03 |
| ATP13A3 | 1.54 | 2.0E-13 |
| OASL | 1.53 | 2.8E-02 |
| TRIM36 | 1.53 | 1.0E-06 |
| PCNX2 | 1.53 | 3.4E-05 |
| KCNN4 | 1.53 | 6.3E-04 |
| ODC1 | 1.53 | 8.6E-05 |
| SPA17 | 1.53 | 3.8E-02 |
| MTHFD2 | 1.53 | 5.1E-13 |
| ST3GAL4 | 1.53 | 1.2E-04 |
| PEA15 | 1.52 | 3.9E-08 |
| IFIT3 | 1.52 | 2.7E-02 |
| BCAR1 | 1.52 | 3.5E-02 |
| LAMP3 | 1.51 | 3.6E-02 |
| ANG | 1.51 | 1.2E-04 |
| SLC4A11 | 1.51 | 5.4E-02 |
| POLR1G | 1.51 | 3.4E-03 |
| HOXC4 | 1.51 | 6.1E-03 |
| NDRG1 | 1.51 | 5.0E-14 |
| TSC22D1 | 1.51 | 2.8E-05 |
| DUSP14 | 1.50 | 2.8E-03 |
| TNNI2 | 1.50 | 9.6E-04 |
| ITGAX | 1.50 | 4.2E-12 |
| SEMA4C | 1.50 | 5.4E-05 |
| CCR5 | 1.50 | 2.5E-05 |

### Supplementary Table 4_continued

**b** **Up in FOLR2 Mac**: 341 genes

| Gene | log2FC | padj |
|---|---|---|
| CCDC141 | -6.61 | 1.0E-35 |
| TTN | -6.52 | 1.4E-28 |
| IL12RB2 | -6.49 | 7.0E-25 |
| SCN9A | -6.35 | 7.0E-37 |
| **LYVE1** | **-6.11** | **1.6E-29** |
| PLEKHG5 | -6.00 | 1.0E-27 |
| P2RY14 | -5.76 | 1.0E-24 |
| ABCA9 | -5.61 | 1.7E-29 |
| ABCA6 | -5.36 | 1.5E-48 |
| C4BPB | -5.25 | 9.7E-14 |
| MAMDC2 | -5.17 | 8.5E-30 |
| CLEC4G | -5.14 | 2.0E-17 |
| TNFRSF25 | -5.07 | 2.1E-34 |
| CCL13 | -4.93 | 1.1E-17 |
| FCER2 | -4.86 | 3.6E-23 |
| SCML2 | -4.53 | 2.9E-10 |
| KHDRBS2 | -4.49 | 4.0E-11 |
| CD163L1 | -4.43 | 4.0E-56 |
| SLC39A12 | -4.36 | 6.2E-08 |
| MPPED2 | -4.31 | 3.5E-23 |
| DNM1 | -4.24 | 8.9E-22 |
| EGFL7 | -4.11 | 2.3E-31 |
| GRIN2C | -4.08 | 3.1E-06 |
| TRIM50 | -4.08 | 1.6E-17 |
| F13A1 | -4.04 | 6.1E-36 |
| COCH | -4.01 | 2.9E-03 |
| DAAM2 | -3.98 | 2.4E-06 |
| LGI2 | -3.88 | 9.8E-21 |
| GFRA2 | -3.87 | 4.9E-31 |
| RPH3AL | -3.85 | 5.6E-20 |
| HRH1 | -3.80 | 2.0E-48 |
| F2RL3 | -3.77 | 8.4E-04 |
| KL | -3.70 | 1.1E-06 |
| PCDH12 | -3.68 | 2.1E-15 |
| CD209 | -3.68 | 1.1E-41 |
| OR2L3 | -3.64 | 7.5E-04 |
| KBTBD12 | -3.61 | 3.1E-05 |
| LILRB5 | -3.58 | 3.6E-53 |
| CFP | -3.44 | 9.6E-10 |
| SLITRK4 | -3.37 | 8.1E-09 |
| EMID1 | -3.34 | 1.2E-08 |
| CCL8 | -3.28 | 8.9E-07 |
| SIGLEC11 | -3.26 | 9.2E-18 |
| **MRC1** | **-3.26** | **6.3E-30** |
| ARL5C | -3.26 | 2.2E-11 |
| TMEM236 | -3.25 | 2.8E-17 |
| OR2L5 | -3.19 | 1.6E-04 |
| KCNA5 | -3.13 | 8.8E-03 |
| PID1 | -3.11 | 1.1E-27 |
| BMP2 | -3.10 | 2.9E-07 |
| GIPC2 | -3.08 | 5.7E-05 |
| TDRD10 | -3.04 | 1.6E-05 |
| SHE | -3.02 | 5.8E-10 |
| TMIE | -2.99 | 2.4E-05 |
| PKD1L3 | -2.99 | 3.1E-13 |
| GARIN4 | -2.97 | 8.7E-03 |
| LRRC4 | -2.95 | 7.8E-09 |

| Gene | log2FC | padj |
|---|---|---|
| CLEC3B | -2.95 | 7.2E-06 |
| ABCD2 | -2.93 | 1.1E-05 |
| SYT1 | -2.93 | 1.3E-03 |
| SLC16A7 | -2.93 | 3.5E-17 |
| KANK2 | -2.92 | 1.1E-13 |
| ALS2CL | -2.90 | 5.3E-14 |
| COLEC12 | -2.87 | 6.8E-23 |
| FZD9 | -2.87 | 1.3E-02 |
| LMO1 | -2.82 | 2.4E-02 |
| METTL27 | -2.79 | 9.5E-15 |
| EDA | -2.74 | 1.7E-17 |
| NLRP6 | -2.71 | 3.4E-03 |
| PPM1N | -2.71 | 3.6E-04 |
| KCNG2 | -2.71 | 7.2E-04 |
| NRXN2 | -2.71 | 2.1E-07 |
| IQCD | -2.70 | 5.6E-09 |
| FGF13 | -2.70 | 9.5E-06 |
| MTSS1 | -2.70 | 5.2E-52 |
| FXYD2 | -2.69 | 4.5E-08 |
| WLS | -2.69 | 1.6E-20 |
| GULP1 | -2.68 | 8.1E-04 |
| STAB1 | -2.68 | 6.1E-26 |
| GPR171 | -2.66 | 3.2E-03 |
| RCN3 | -2.66 | 7.0E-33 |
| KLF2 | -2.66 | 2.3E-21 |
| PDE4D | -2.65 | 1.7E-11 |
| **FOLR2** | **-2.64** | **9.1E-38** |
| ACTN2 | -2.64 | 1.1E-02 |
| RGL1 | -2.64 | 3.3E-44 |
| IRAG1 | -2.62 | 3.9E-14 |
| ASB9 | -2.57 | 3.8E-07 |
| TNFSF18 | -2.57 | 2.8E-04 |
| CBFA2T3 | -2.56 | 6.2E-10 |
| ITSN1 | -2.56 | 3.2E-40 |
| GPBAR1 | -2.56 | 6.5E-04 |
| **WWP1** | **-2.55** | **3.4E-47** |
| PGA5 | -2.53 | 1.0E-01 |
| EFHC2 | -2.53 | 4.0E-02 |
| **CETP** | **-2.52** | **1.9E-02** |
| RNASE1 | -2.52 | 1.7E-06 |
| NEURL2 | -2.52 | 6.9E-07 |
| SPIC | -2.52 | 1.6E-02 |
| FOSB | -2.51 | 8.3E-08 |
| CD200R1 | -2.50 | 5.7E-27 |
| CLEC10A | -2.50 | 1.5E-19 |
| AFF3 | -2.47 | 8.3E-14 |
| FIGNL2 | -2.46 | 2.2E-02 |
| IL2RA | -2.45 | 9.8E-09 |
| CCL2 | -2.45 | 7.0E-05 |
| OSBPL6 | -2.44 | 1.1E-04 |
| TEKT3 | -2.43 | 5.7E-02 |
| GPER1 | -2.42 | 1.4E-03 |
| NEU4 | -2.42 | 1.0E-02 |
| ENSG00000290796 | -2.41 | 2.0E-03 |
| OR2AK2 | -2.41 | 2.8E-02 |
| ARHGEF37 | -2.41 | 9.5E-07 |
| IL7 | -2.40 | 8.1E-16 |

| Gene | log2FC | padj |
|---|---|---|
| TSPAN7 | -2.39 | 1.8E-03 |
| MARCO | -2.38 | 2.8E-07 |
| GAS6 | -2.37 | 7.6E-20 |
| RGL3 | -2.36 | 4.3E-15 |
| PLEKHG3 | -2.36 | 7.1E-19 |
| KIAA1549 | -2.36 | 5.5E-10 |
| DAB2 | -2.36 | 3.1E-56 |
| KLF4 | -2.35 | 7.4E-09 |
| ENHO | -2.35 | 5.6E-02 |
| CCDC89 | -2.35 | 3.4E-03 |
| SOX5 | -2.35 | 6.3E-04 |
| PCDHGA9 | -2.33 | 2.0E-03 |
| TNFRSF4 | -2.32 | 2.5E-03 |
| PRR5 | -2.31 | 2.1E-12 |
| DIRAS3 | -2.31 | 8.0E-06 |
| CCL14 | -2.30 | 3.0E-06 |
| HPN | -2.30 | 6.2E-07 |
| SCN1B | -2.28 | 2.8E-12 |
| SLC24A4 | -2.27 | 2.2E-03 |
| MDGA1 | -2.27 | 6.9E-02 |
| DCHS1 | -2.27 | 8.4E-16 |
| CPAMD8 | -2.24 | 5.2E-04 |
| IGF1 | -2.23 | 1.7E-36 |
| STON2 | -2.23 | 3.5E-15 |
| NPAS3 | -2.21 | 2.3E-03 |
| ATOH8 | -2.21 | 5.4E-03 |
| CXCL12 | -2.20 | 4.6E-15 |
| PRKCQ | -2.19 | 2.3E-02 |
| HSF4 | -2.17 | 7.1E-08 |
| SLC9A9 | -2.17 | 1.2E-44 |
| CR1 | -2.17 | 1.4E-16 |
| SPTBN5 | -2.15 | 2.9E-04 |
| SKP2 | -2.15 | 1.7E-18 |
| VIPR1 | -2.15 | 5.4E-04 |
| GALNT14 | -2.14 | 4.6E-02 |
| CES3 | -2.13 | 1.5E-05 |
| CD28 | -2.12 | 4.2E-09 |
| MPP3 | -2.12 | 4.9E-06 |
| DMWD | -2.12 | 6.2E-08 |
| PLTP | -2.11 | 1.0E-11 |
| SKIDA1 | -2.11 | 5.2E-02 |
| ROBO4 | -2.11 | 2.1E-05 |
| ARHGAP8 | -2.11 | 2.0E-06 |
| SELENOP | -2.11 | 1.6E-11 |
| RIBC2 | -2.10 | 1.1E-02 |
| C11orf91 | -2.09 | 4.0E-02 |
| NXPH3 | -2.09 | 6.0E-03 |
| SLC43A1 | -2.09 | 9.7E-07 |
| PYROXD2 | -2.08 | 1.7E-09 |
| NR3C2 | -2.07 | 2.6E-09 |
| LHFPL6 | -2.07 | 1.7E-06 |
| TSLP | -2.07 | 5.4E-02 |
| MNS1 | -2.06 | 7.5E-05 |
| TESC | -2.05 | 1.2E-03 |
| TOMM20L | -2.05 | 1.1E-01 |
| FGFR1 | -2.05 | 2.6E-11 |
| PABPN1L | -2.04 | 8.3E-02 |

| Gene | log2FC | padj |
|---|---|---|
| TSPYL2 | -2.03 | 1.1E-12 |
| NOL3 | -2.03 | 3.5E-07 |
| RNF150 | -2.03 | 1.1E-04 |
| WDR97 | -2.03 | 1.7E-05 |
| SLC15A2 | -2.02 | 8.0E-14 |
| SMO | -2.02 | 1.2E-05 |
| DDX43 | -2.02 | 9.2E-02 |
| SIGLEC1 | -2.01 | 1.6E-15 |
| BAHCC1 | -2.00 | 6.5E-10 |
| SHROOM1 | -2.00 | 2.2E-07 |
| RIMKLB | -2.00 | 7.6E-13 |
| MACIR | -2.00 | 1.1E-04 |
| PDE7B | -2.00 | 6.5E-02 |
| SGMS1 | -1.99 | 1.4E-54 |
| SHC2 | -1.99 | 3.6E-03 |
| ANKUB1 | -1.99 | 2.9E-02 |
| STARD13 | -1.98 | 1.3E-16 |
| NYNRIN | -1.98 | 7.2E-08 |
| **VCAM1** | **-1.98** | **8.8E-05** |
| CNRIP1 | -1.97 | 6.4E-24 |
| ARHGAP23 | -1.96 | 9.2E-04 |
| SLC22A23 | -1.95 | 7.0E-11 |
| IGFBP4 | -1.95 | 1.1E-11 |
| LEFTY1 | -1.95 | 2.3E-02 |
| RSPH4A | -1.95 | 8.3E-02 |
| IQSEC3 | -1.95 | 6.1E-02 |
| SHMT1 | -1.94 | 2.2E-22 |
| NEXMIF | -1.94 | 9.7E-02 |
| NSUN7 | -1.93 | 5.5E-06 |
| DMPK | -1.93 | 2.0E-03 |
| PDE2A | -1.93 | 3.6E-03 |
| HTR1F | -1.91 | 2.8E-02 |
| CSGALNACT1 | -1.91 | 1.6E-05 |
| LRP5 | -1.90 | 1.4E-04 |
| TSHZ3 | -1.90 | 5.1E-13 |
| PRSS36 | -1.89 | 3.9E-13 |
| CCDC183 | -1.89 | 2.1E-02 |
| SCAMP5 | -1.88 | 6.4E-09 |
| THBD | -1.88 | 6.0E-08 |
| ME1 | -1.88 | 9.9E-26 |
| HIP1 | -1.88 | 1.0E-08 |
| FAM20A | -1.87 | 4.5E-17 |
| CHADL | -1.87 | 1.5E-02 |
| KIAA1614 | -1.87 | 1.4E-05 |
| WDFY2 | -1.86 | 7.3E-23 |
| PEAK1 | -1.86 | 3.0E-17 |
| BIRC3 | -1.85 | 5.7E-07 |
| DYNLT4 | -1.85 | 2.4E-02 |
| HCN2 | -1.85 | 1.2E-02 |
| LTBP2 | -1.85 | 1.5E-07 |
| MAF | -1.84 | 4.8E-40 |
| CUZD1 | -1.84 | 4.6E-02 |
| SERPINI2 | -1.84 | 1.1E-01 |
| SULT1C4 | -1.84 | 3.3E-03 |
| ADAM33 | -1.83 | 5.9E-03 |
| ADAMTS14 | -1.83 | 5.8E-03 |
| HOXB6 | -1.83 | 4.0E-03 |

| Gene | log2FC | padj |
|---|---|---|
| DCST2 | -1.82 | 3.2E-02 |
| DLGAP3 | -1.82 | 1.0E-02 |
| GNG2 | -1.81 | 3.0E-14 |
| ST8SIA1 | -1.81 | 3.8E-02 |
| PRRT2 | -1.80 | 6.2E-02 |
| CXCL3 | -1.80 | 7.1E-03 |
| FAXDC2 | -1.80 | 3.8E-13 |
| INKA1 | -1.80 | 3.4E-06 |
| GJC2 | -1.79 | 1.3E-02 |
| PMAIP1 | -1.79 | 3.2E-06 |
| SLIT1 | -1.78 | 5.3E-02 |
| KITLG | -1.78 | 2.3E-02 |
| KATNAL2 | -1.77 | 2.4E-06 |
| NRP1 | -1.76 | 5.4E-15 |
| ARMC2 | -1.76 | 8.3E-07 |
| KRBA1 | -1.75 | 1.8E-06 |
| ABLIM2 | -1.74 | 1.4E-02 |
| IGHM | -1.73 | 5.2E-02 |
| **CD163** | **-1.73** | **1.2E-15** |
| MMP25 | -1.73 | 1.3E-02 |
| CTRC | -1.72 | 2.9E-02 |
| PCDHGA11 | -1.72 | 2.2E-03 |
| NAALAD2 | -1.71 | 9.6E-02 |
| CXCL1 | -1.71 | 1.1E-01 |
| SCARA5 | -1.71 | 3.3E-02 |
| C3orf70 | -1.71 | 2.1E-02 |
| RHOJ | -1.70 | 5.4E-02 |
| STARD8 | -1.70 | 6.6E-08 |
| OLFML1 | -1.70 | 7.9E-02 |
| SULT1A2 | -1.70 | 8.3E-02 |
| DPF3 | -1.70 | 4.4E-02 |
| ADAM11 | -1.69 | 4.2E-02 |
| S1PR1 | -1.69 | 2.2E-05 |
| NLRP3 | -1.69 | 2.1E-05 |
| PIWIL2 | -1.69 | 9.7E-02 |
| DENND2A | -1.69 | 3.4E-02 |
| JPH4 | -1.69 | 1.5E-03 |
| GPSM1 | -1.68 | 1.5E-05 |
| PSPN | -1.68 | 4.5E-02 |
| TBC1D14 | -1.68 | 2.2E-26 |
| NXF3 | -1.68 | 2.4E-02 |
| GVQW3 | -1.68 | 6.3E-04 |
| ARHGAP6 | -1.68 | 1.9E-07 |
| **SLC40A1** | **-1.68** | **7.2E-16** |
| C5AR2 | -1.68 | 8.9E-09 |
| DMD | -1.67 | 2.1E-08 |
| TPPP3 | -1.67 | 7.9E-03 |
| MS4A4A | -1.67 | 1.5E-15 |
| FRMD4B | -1.66 | 3.9E-24 |
| SOCS3 | -1.66 | 6.6E-06 |
| VEGFD | -1.66 | 8.2E-02 |
| ADGRA2 | -1.65 | 1.7E-04 |
| UNC5CL | -1.65 | 1.0E-02 |
| GABRB2 | -1.65 | 8.2E-02 |
| P2RY6 | -1.65 | 1.1E-09 |
| ZNF324B | -1.65 | 1.9E-02 |
| ZBTB16 | -1.64 | 3.7E-02 |

| Gene | log2FC | padj |
|---|---|---|
| MMP17 | -1.64 | 1.3E-02 |
| FAM124B | -1.64 | 1.8E-02 |
| FAM118A | -1.64 | 8.1E-04 |
| PCDHGB7 | -1.63 | 8.7E-09 |
| PLEKHH2 | -1.63 | 7.5E-03 |
| PMP22 | -1.62 | 6.9E-10 |
| GLIS3 | -1.62 | 1.3E-04 |
| ADAMTS2 | -1.62 | 5.3E-03 |
| PRDM8 | -1.62 | 7.8E-02 |
| IQCN | -1.62 | 2.0E-06 |
| NEIL1 | -1.62 | 3.9E-06 |
| EMB | -1.61 | 8.0E-09 |
| EGFR | -1.61 | 1.4E-02 |
| ALDH8A1 | -1.61 | 1.8E-02 |
| IGLC1 | -1.61 | 3.4E-02 |
| GPRC5B | -1.61 | 2.4E-05 |
| EGFL8 | -1.59 | 1.4E-05 |
| TEDC1 | -1.59 | 6.3E-04 |
| FCAR | -1.59 | 2.3E-03 |
| SLC41A1 | -1.59 | 1.1E-04 |
| CXCL2 | -1.59 | 5.8E-04 |
| GCNT1 | -1.58 | 3.1E-12 |
| ZNF117 | -1.58 | 6.0E-08 |
| ARHGEF33 | -1.58 | 2.7E-02 |
| SLC25A27 | -1.58 | 5.2E-02 |
| **ENPP2** | **-1.57** | **5.1E-05** |
| PDGFC | -1.57 | 2.2E-26 |
| VSIG4 | -1.57 | 1.3E-09 |
| MAN1A1 | -1.56 | 2.9E-13 |
| TLE4 | -1.56 | 2.7E-11 |
| NID1 | -1.55 | 4.8E-04 |
| WDR17 | -1.55 | 2.1E-05 |
| HTR2B | -1.55 | 6.4E-03 |
| FOXRED2 | -1.55 | 3.5E-09 |
| LDLR | -1.55 | 4.1E-03 |
| MPEG1 | -1.55 | 4.1E-23 |
| FBXO24 | -1.55 | 7.9E-02 |
| DNAAF9 | -1.54 | 6.7E-21 |
| GPR162 | -1.54 | 1.1E-08 |
| RSPO4 | -1.54 | 1.0E-01 |
| CCDC170 | -1.54 | 1.4E-10 |
| FGF7 | -1.54 | 1.9E-02 |
| KREMEN1 | -1.53 | 2.8E-03 |
| GARNL3 | -1.53 | 4.4E-04 |
| MAPK12 | -1.53 | 5.4E-04 |
| OLFM1 | -1.53 | 3.7E-02 |
| NCF4 | -1.52 | 7.6E-30 |
| PTGIR | -1.52 | 4.2E-03 |
| FAM13A | -1.52 | 2.3E-12 |
| HOXA10 | -1.52 | 8.4E-02 |
| EPS15 | -1.52 | 1.8E-27 |
| FOS | -1.52 | 1.6E-13 |
| RHOBTB1 | -1.51 | 1.5E-23 |
| RDH16 | -1.51 | 8.4E-03 |
| EFCAB2 | -1.50 | 1.0E-05 |
| EGR1 | -1.50 | 3.3E-04 |

**Supplementary Table 4 Differentially expressed genes between TREM2⁺ Macs and FOLR2⁺ Macs. a.** Results of differential gene expression analysis in the DCIS snRNA-seq dataset. Genes with higher expression in TREM2⁺ macrophages (logFC > 1.5, adjusted P < 0.01) are shown. Genes previously reported as differentially expressed in the respective macrophage subsets in breast cancer are highlighted in red. **b.** Genes with higher expression in FOLR2⁺ macro-phages (logFC > 1.5, adjusted P < 0.01) are shown.

#### Supplementary Table 5

| Antigen | Clone | Provider | Catalogue number | Dilution | Antigen retrival buffer | Fluorephore |
|---|---|---|---|---|---|---|
| Pan-CK | AE-1/AE-3 | Novus | NBP2-29429 | 1:800 | ER1 | OPAL 780 |
| αSMA | 1A4 | Invitrogen | 14-9760-82 | 1:400 | ER2 | OPAL 480 |
| IBA1 | EPR6136(2) | abcam | ab178680 | 1:500 | ER2 | OPAL 570 |
| TREM2 | D8I4C | Cell Signaling Tech | 91068 | 1:100 | ER2 | OPAL 690 |
| FOLR2 | OTI4G6 | Invitrogen | MA5-26933 | 1:150 | ER2 | OPAL 620 |
| PGRN | 2D4-2F1 | Invitrogen | MA1187 | 1:100 | ER2 | OPAL 520 |
| SPP1 | RM1018 | abcam | ab283656 | 1:500 | ER2 | OPAL 520 |
| GAL9 | RM499 | Invitrogen | MA5-49315 | 1:100 | ER2 | OPAL 520 |

**Supplementary Table 5 Primary antibodies used for multiplex fluorescence immunohistochemistry.** CK, cytokeratin; αSMA, alpha smooth muscle actin; IBA1, Ionized calcium-binding adapter molecule 1; TREM2, triggering receptor expressed on myeloid cells 2; FOLR2, folate receptor 2; PGRN, progranulin; SPP1, secreted phosphoprotein 1; GAL9, galectin 9; ER1, BOND Epitope Retrieval Buffer 1; ER2, BOND Epitope Retrieval Buffer 2.
